# Mapping Coastal Forest Retreat using Convolutional Neural Networks and Different Satellite Imagery

**DOI:** 10.64898/2026.08.18.745552

**Authors:** Titilayo T. Tajudeen, Marcelo Ardon, Mirela Tulbure, Katherine L. Martin

## Abstract

Coastal forests are increasingly threatened by saturated soil and elevated salinity levels resulting from sea level rise, saltwater intrusion, and storm surges. In response to rising salinization and flooding, healthy coastal forests that rely on freshwater (both wetland forests and low-elevation upland forests) are transitioning into landscapes dominated by dead or dying trees, known as ghost forests. Situated among salt-tolerant shrubs and grasses, ghost forests eventually become marshes or open water. Here, our main objective was to quantify the dynamics and pathways of these forest landscape conversions, as well as the factors contributing to the changes, which is vital for understanding the progression of coastal ecosystem degradation and forecasting future changes. We focused first on identifying the best method to track forest landscape change by exploring the role of multiple remote sensing indices (i.e., multispectral, bi-seasonal, topographical, and phenological metrics) in enhancing the performance of deep learning models (convolutional neural networks, CNNs) for land cover classification in the coastal plain of North Carolina using surface reflectance of Landsat 8 and Sentinel-2 images. Then, we used the best available data (Landsat 8) to understand long-term change and identify patterns of land cover change from 1985 to 2021. Our study reveals that incorporating phenology and topographical indices enhances the separability of the ghost forests class from all other vegetation classes. In our assessment, the higher-resolution Sentinel-2 data (F1 Score = 96.3) outperformed Landsat images (F1 score = 93.4) for the 2021 co-available year. However, Landsat remains an important tool used due to its long-term data record. Therefore, we used Landsat to determine that 21% of forests were lost between 1985 and 2021, and that the rate of loss is increasing. Between 2010 and 2021, 23,876 ha of forest were converted to marsh, ghost forest, and shrub, which is 1.5 times higher than the 16,968 ha lost between 1985 and 2010. These conversions from forest to ghost forest and marshes were driven primarily by proximity to the channel, salinity, and the increasing rate of relative sea level rise (RSLR), which are the key environmental drivers of observed changes. By quantifying these changes, we highlight regions most vulnerable to environmental stressors, providing a basis for targeted conservation strategies.

## 1 Introduction

Ghost forests in coastal areas, characterized by the presence of dead or dying trees, are striking indicators of sea level rise, increased salinity, and climate change (1–4). Along the edges of the mid-Atlantic and low-lying areas of the US East Coast, rising sea levels are forcing saltwater and waterlogging further inland, altering forest ecosystems (5). The coastal plain along the U.S. Atlantic coast is a recognized global hotspot for rising sea levels (6) spanning through a strong gradient of salinity, topography, and a high rate of relative sea-level rise (RSLR) (1). White et al. (2022) (7) estimated that over 13,600 km^2^ (i.e., 1,360,000 ha, 8%) of forested wetlands transitioned to ghost forests, shrub-scrub, and/or marshes along the coastal plain of the eastern US coast between 1996 and 2016. This loss rate is higher than global estimates of mangrove loss(7). Current rates of coastal forest lateral retreat, a landward shift, are 2-14 times faster than pre-industrial rates (2,8). Within the Alligator River National Wildlife Refuge, located in the Albemarle-Pamlico Peninsula of North Carolina, an estimated 19,300 ha of forest (of the 76,575 ha of forests in 1985) transitioned into ghost forests or marshes between 1985 and 2019 (9).

A range of factors contributes to forests shifting toward landscapes of standing dead trees, expanding salt marshes, or open water. Saltwater intrusion and rising sea level jointly have a significant adverse impact on both the function and structure of coastal forests (4,10–12). Persistent stress from soil salinization and rising inundation causes forest dieback and a long-term shift in coastal plant communities (13). The eastern US Coastal Plains are also characterized by low topographic relief, further increasing the vulnerability of these ecosystems (5,14).

Exposure of forests to salinity increases as marine salts move landward through surface or groundwater connections in artificial canals and ditches originally installed to reduce inundation (15–17). Coastal forests are therefore becoming more vulnerable as artificial drainage systems serve as pathways for saltwater intrusion during droughts and coastal storms, when storm surges or high tides push brackish water inland (18,19). Artificial drainage systems along North Carolina’s eastern coast increase the drainage density in the landscape, reduce flow accumulation that can push out saltwater, making low elevation more vulnerable to saltwater intrusion (15).

High-resolution mapping and monitoring of ghost forests is vital for understanding the progression and pathways of coastal ecosystem loss and transition, informing conservation efforts, and predicting future changes (20). Satellite-based remote sensing has emerged as a vital tool for this purpose, providing large-scale and long-term data that can capture the dynamic processes leading to ecosystem transition and loss (21–24). While previous studies have employed remote sensing approaches to map ghost forests (4,7,9), these studies mainly relied on using 30-meter Landsat data. Hence, several advances may improve upon current estimates of the rate and drivers of coastal forest loss by using higher-resolution data in combination with other metrics.

In particular, land surface phenological (LSP) metrics are usually linked to overall inter-annual vegetation changes that can be understood from spectral remote sensing imagery. These include the start of the greening or growing season (SOS), the peak of the growing season (POS), the onset of senescence, the length of the growing season, and the end of the season (EOS) (25,26). A major challenge associated with classifying land cover over time is that the spatial and spectral characteristics of a single class change throughout the year (27). The difficulty increases when monitoring across multiple years, as differences in phenological conditions, which are unpredictable from year to year, also affect model predictions (24). These shifts and inconsistencies in phenology have been linked to shifts in the timing and intensity of seasons caused by climate change, which in turn affect the reflectance values captured by sensors (28–30). Although focusing on a single moment in time can be confusing for the model, the intra-annual phenological change may hold key information for differentiating vegetation classes, provided that these patterns can be integrated into the model (24,31). For instance, phenology in ghost forests may be distinctive because, unlike healthy vegetation that follows seasonal patterns of greening and senescence, ghost forests exhibit reduced or absent vegetative activity, characterized by dead trees, which can alter the typical seasonal signals captured by remote sensing (32). Similarly, phenological changes could help distinguish between vegetation classes, such as shrubs (9) and healthy or degraded classes due to spectral similarity.

Furthermore, machine learning approaches such as decision trees (DTs) (7) and random forest (RF) (7,32,33) have been used for ghost forest identification from satellite images, while support vector machines (SVMs) and other traditional classifiers have been used in land cover mapping. SVM received considerable attention because of its ability to handle high-dimensional data and its effectiveness in performing well with limited training samples (34). Meanwhile, RF became popular for its ease of use, such as being relatively insensitive to classification parameters and typically achieving high accuracy (35). Although land cover classification has long been a central focus of remote sensing, high intraclass variance (significant differences within a single class) and low inter-class variance (minimal variability between different classes) continue to challenge traditional approaches (36,37), especially the close spectral similarity between ghost forests, forests, and shrubs (9). Deep learning methods represent a promising approach, supported by studies that have demonstrated their ability to surpass conventional models often (38).

The rise of deep learning (DL) has generated interest in using neural networks to classify and analyze remotely sensed data (38). DL algorithms have achieved notable success in various image analysis tasks, such as land use and land cover classification (27,39), data fusion (40), scene classification (41), change detection, and object detection (41). The success is due mainly to the powerful extraction capability of multiple features from annotated samples (36). Deep learning is a learning algorithm based on neural networks (42). Due to their ability to capture spatial hierarchies through local receptive fields and shared weights, Convolutional Neural Networks (CNNs) often outperform traditional one-dimensional models when applied to spatially structured data (43,44).

Previous studies of coastal forest change in North Carolina relied largely only on (a) the spectral indices for summer images, or both summer and winter images (4,9), (b) the use of 30-meter resolution Landsat images only (4,9), and (c) multispectral vegetation indices without the integration of phenological metrics and no application of a deep learning approach. We hypothesized that incorporating phenological indices and/or higher resolution imagery to train a DL model could lead to improved classification accuracy over models trained only on single or bi-seasonal images (27,33,45). Therefore, we focused on developing and applying methods that use satellite imagery at various resolutions and phenological analysis to map transformations in coastal forests along the North Carolina coast. We sought to provide a comprehensive approach to examining changes in these coastal ecosystems by leveraging the strengths of various satellite data sources and phenological indicators. We expected that integrating phenological data with satellite imagery across various resolutions (i.e., 10m and 30m) could enhance the ability to map and monitor ecosystem loss and transition with greater accuracy and temporal sensitivity. Therefore, our objectives were as follows:

a. To evaluate how different satellite resolutions and phenological characteristics can be used to map coastal forests using convolutional neural networks (CNNs) using images taken in 2021.
b. To examine the dynamics of forest change in the study area over a long-term period (1985–2021)
c. To identify the factors driving coastal forest change in the study area over a long-term period (1985–2021)

## 2 Materials and Methods

### 2.1 Study Area

The study area spans 35.28°N to 36.00°N and 75.71°W to 77.12°W along the eastern US Coast, covering 644,892 ha of the Atlantic Coastal Plain in North Carolina, United States (Fig 1). This area includes a series of low-relief drowned river valleys with a chain of barrier islands surrounding the country’s second-largest estuary, the Albemarle-Pamlico. It is right below the US mid-Atlantic coast, an area known as a global sea-level rise hotspot (6), spanning strong gradients in salinity, topography, and accelerating relative sea level rise rate (RSLR). Specifically, this area is experiencing a rapid rate of relative sea-level rise, estimated at 3.5 mm yr^-1^ in 2010 and 5.14 mm yr^-1^ in 2021 (S6 Fig.). Across the study area, hurricanes and SLR are driving saltwater intrusion and increased inundation, resulting in forest dieback (i.e., ghost forest, see Figure 1(a)) and ultimately leading to marsh encroachments (7,9,46,47). The study area encompasses both private and protected areas (48), including the Alligator River National Wildlife Refuge, Pocosin Lakes National Wildlife Refuge, Mattamuskeet National Wildlife Refuge, Buckridge Coastal Reserve, Palmetto Peartree Preserve, and Swanquarter National Wildlife Refuge. Freshwater, shrub-dominated pocosins are unique, acidic, peat-rich wetlands found in the coastal plain, known for their dense vegetation and critical role in water storage (49). Additionally, the eastern part of the study area is mostly covered by woody or forested wetlands; as you move inland, it increasingly consists of upland forests. Farmland also accounts for 27% of the study area.

**Figure 1.**
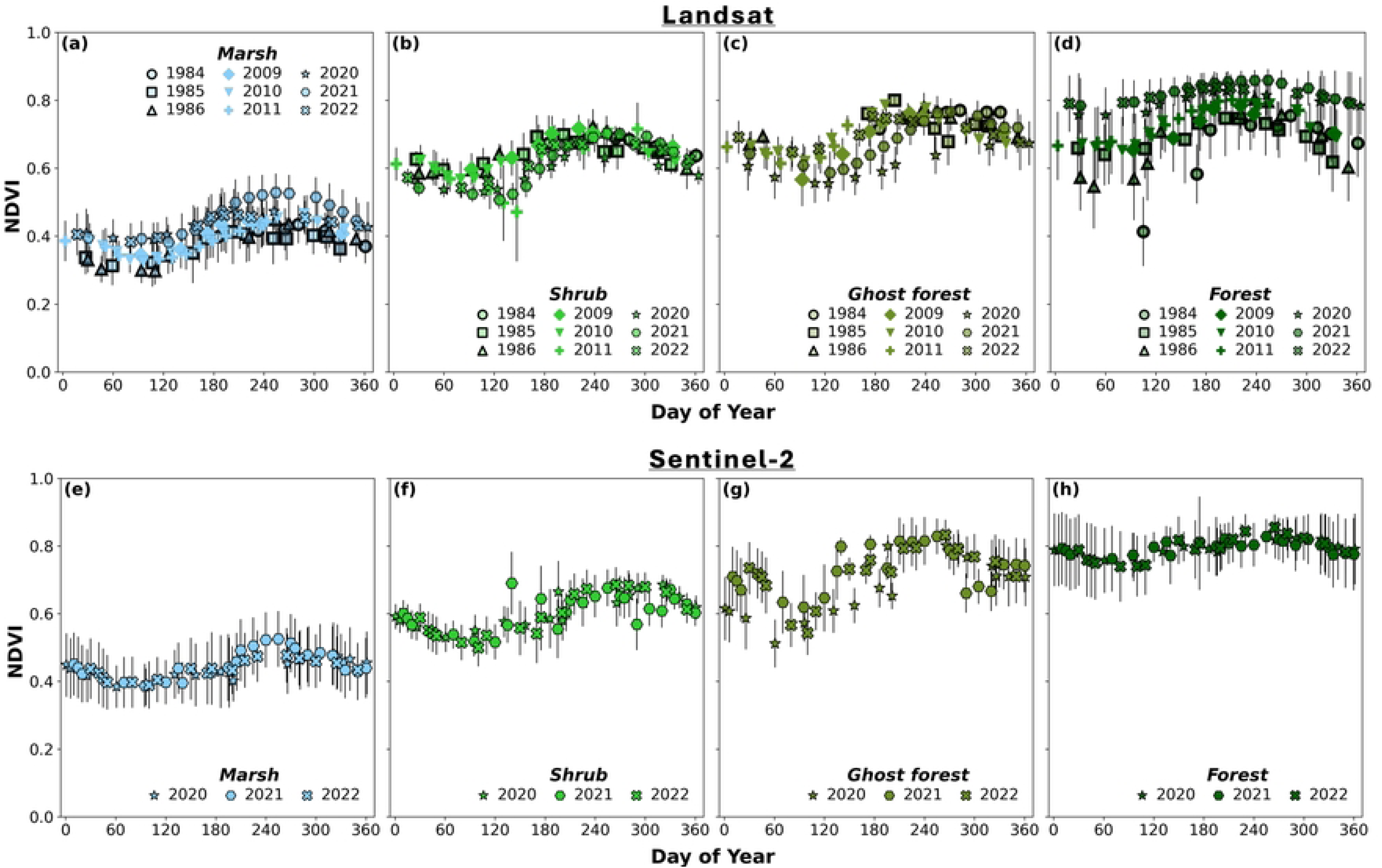
The study area map (bounded in the north by the Albemarle Sound and in the south by the Pamlico Sound) shows the digital elevation model of the North Carolina coastal plain and the protected area (red strip diagonal lines). The image inserts show ghost forests, which increase light availability for shrub growth and marshlands.

### 2.2 Image acquisition and preprocessing

We collected and preprocessed satellite imagery from multiple sensors to classify land cover. We selected all images for the growing and leaf-off seasons from Landsat (30-meter resolution) and Sentinel-2 (10-meter resolution) using a cloud cover filter threshold of less than 2% (see Table 1). The filtered images spanning multiple years (1985, 2010, 2021) and overlapping with the study area were downloaded from the Google Earth Engine catalog. We didn’t account for intermediate time steps before 2010, given the relatively slow processes of coastal forest change, especially since a major portion of forest change in this study area became more visible with the coarse resolution of Landsat images post-2010 (4,9). Images collected included the filtered Landsat 8 and 5 Level 2, Collection 2 Surface Reflectance for six spectral bands (Blue, Green, Red, Near Infrared, Short-wave infrared (SWIR1, SWIR2)) bands, as well as the Harmonized Sentinel-2 Surface Reflectance images (Blue (B2), Green (B3), Red (B4), Near Infrared (B8), SWIR 1 (B11), SWIR(B12). We utilized the ancillary quality assessment datasets to process all Landsat images by masking out pixels related to cloud, cloud shadow, missing data, snow, and ice specific to each satellite sensor. Specifically, residual clouds, shadows, and haze were filtered using the SR_ATMOS_OPACITY (the aerosol optical thickness value in Landsat surface reflectance data, indicating the clarity or haziness of the atmosphere when the image was captured) for Landsat 5 by thresholding to exclude those with values less than 0.3. Conversely, the ’SR_AEROSOL’ band was employed to eliminate pixels labeled as high probability of haze in Landsat 8 (50). Images were also visually verified to ensure they were not inundated and of high quality. To be consistent, two images (representing bi-season) were acquired to generate the annual land cover classification map: one in May-September (growing season) and the other in December–February (leaf off) – the period when inter-vegetation contrast is at its highest (9,23,32,51). The 2019 and 2011 images were used to supplement the validation and testing data, given our small study area.

**Table 1:**
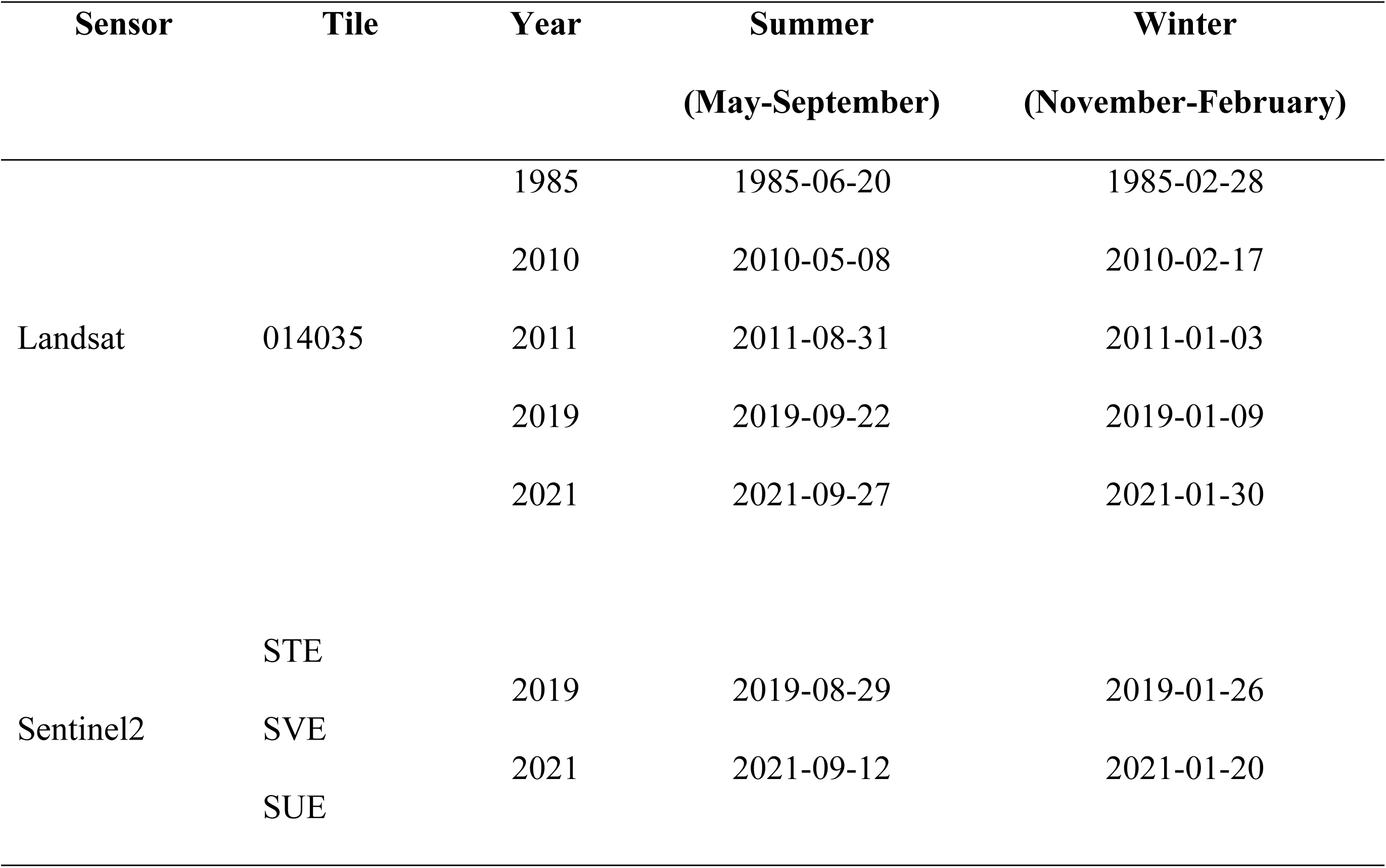
Data acquisition description for the winter and summer seasons.

Multispectral indices, including the normalized difference vegetation index (NDVI), Enhanced Vegetation Index (EVI), modified normalized difference water index (mNDWI), tasseled cap transformation, and Modified Soil-Adjusted Vegetation Index (mSAVI), were calculated and stacked over the spectral bands (see Table 2).

**Table 2.** Summary of metrics from the Landsat 5, 8, and Sentinel 2 images.

| Features | Variables | Description/Formula |
| --- | --- | --- |
| Spectral Bands | Blue, Green, Red, | Surface Reflectance of the blue band, green band, red band, near |
|  | NIR, SWIR1, and | infrared band, short-wave infrared band 1, and short-wave infrared |
|  | SWIR2 | band 2 for the summer (S) and winter (W) seasons |
| Multispectral Indices | NDVI | $NDVI = \frac{NIR - RED}{NIR + RED}$ |
| | EVI | $EVI = \frac{2.5 * (NIR - RED)}{NIR + 6 * RED - 7.5 * BLUE + 1}$ |
| | MSAVI | $MSAVI = 0.5(2 * NIR + 1 \sqrt{(2 * NIR + 1)^2 - 8 * (NIR - RED)})$ |
| | mNDWI | $mNDWI = \frac{GREEN - SWIR1}{GREEN + SWIR1} ((4,32,45,62))$ |
|  | TCP- Wetness | Tasseled Cap Transformation Wetness and Greenness Index computed |
|  | TCP- Greenness | from the surface reflectance of all six bands. Coefficient differ between Landsat 5 (63), Landsat 8 (64), and Sentinel-2 (65) |
| Biophysical Indices | Elevation | Resampled to 30m resolution using bilinear interpolation from the 1m |
|  | Slope | Coastal National Elevation Database (CoNED) DEM (66). |
| Phenological Metrics | SOGS | Start of Growing Seasons |
|  | POGS | Peak of Growing Seasons |
|  | LOGS | Length of Growing Season |
|  | A_NDVI | Amplitude of NDVI |
|  | B_NDVI | Base NDVI |
|  | P_NDVI | Peak NDVI |

To collect training samples for the stacked bands, Esri and Google Earth imagery, collected on March 8, 2021, and April 30, 2010, were used as a reference for visual inspection. Additionally, the NC Orthophoto imagery (6 inches) for the years 2020 and 2010 (52) and the 2019 NAIP Imagery were also used for visual reference (53). The classes included forests, ghost forests, shrubs, marshes, water, and cultivated land. Samples were taken for each class for the years 2021, 2011, and 2010 (see Fig. 2 for a description of the classes). We obtained additional training data for the shrub and ghost forest classes from previous research conducted in the Albemarle Peninsula (7,9). Specifically, as shown in Figure 2 (used for training data), each point characterized as a ghost forest contained approximately 20–40 visible snags or fallen tree trunks per Landsat pixel, with some bare-soil reflectance (9). The forest here refers to upland forests and forested wetlands (i.e., freshwater swamps) with closed-canopy covers.

**Figure 2.**
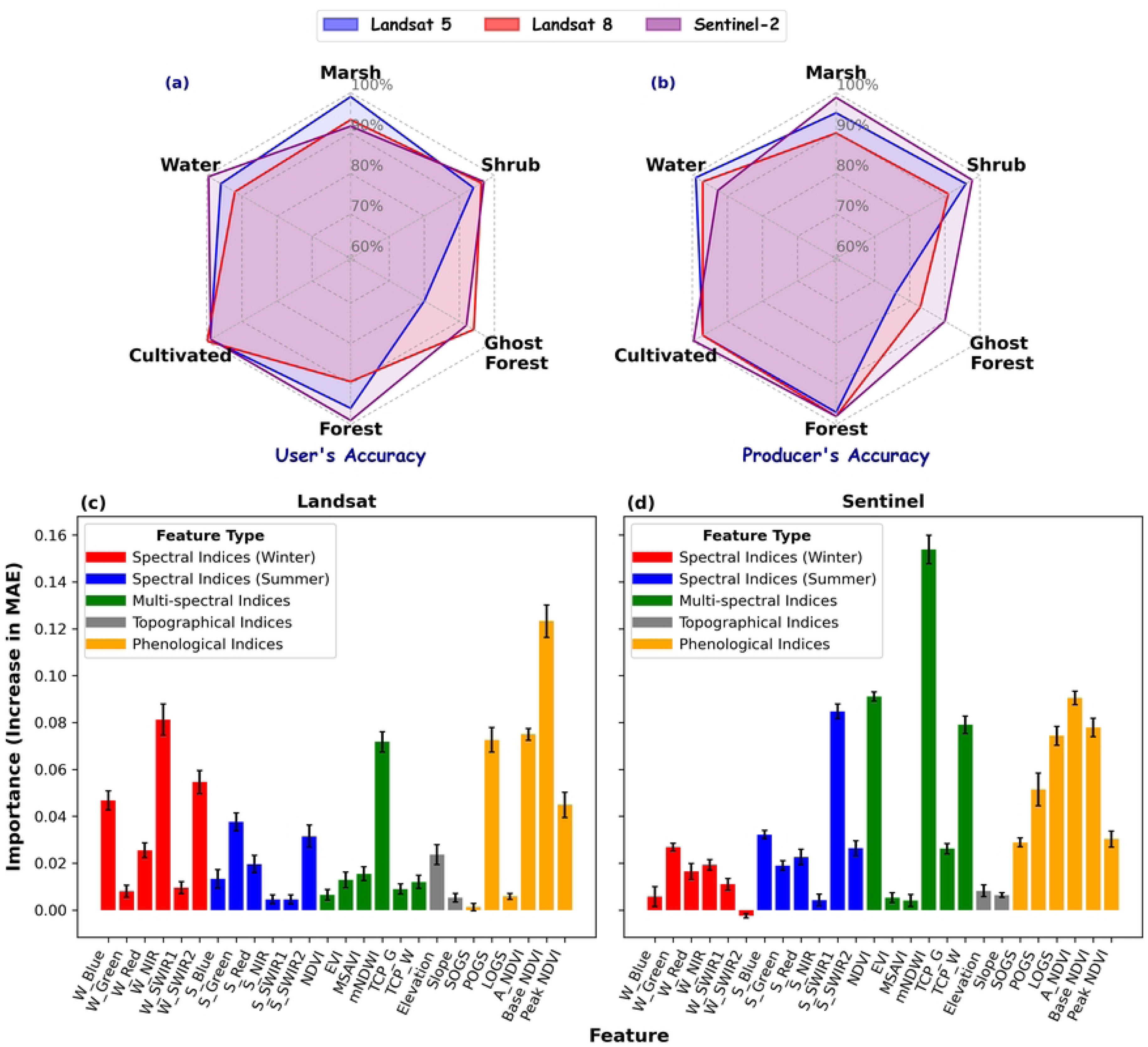
Examples of land cover classes mapped and visualized in a false color image (NIR, Red, and Green). Source: NC Orthophoto Imagery, NC OneMap (2020).

### 2.3 Vegetation phenology and phenological metrics

As many specific phenological events cannot be directly observed at the spatial resolution of satellite imagery, vegetation dynamics are characterized using a descriptor called “land surface phenology (LSP)” (54), which is estimated based on time series vegetation indices such as NDVI derived from satellite data (55). We extracted phenological metrics from the time-series NDVI computed from satellite imagery, after smoothing using the Savitsky-Golay filter (56) to reduce uncertainties in the phenological signals. The Savitsky-Golay function has been widely used to reduce noise in the vegetation index (57,58). The model used for this is Phenolopy (59), a pixel-based phenological extraction Python library. This study adopted the Chen and Kirwan (2022) (32) method, which involves stacking a 3-year NDVI image to account for prominent cloud cover. Firstly, we download Landsat surface reflectance images from Google Earth Engine by filtering to select only images with less than 60% cloud cover and 35% cloud cover for the Sentinel images. These images include both winter and summer images, which are used for classification. Then, we computed a 3-year NDVI (e.g., the previous year – 2020, the ***year of interest – 2021***, and the following year - 2022) for all the masked images (i.e., excluding water, haze, clouds, and cloud shadow). We further stacked all NDVI data in Day of Year (DOY) to produce enough monthly data points for modeling yearly phenological metrics (32). On average across all 3-year NDVI stacks, we have 36 ± 5 images per 3-year for Landsat and 51 ± 9 images per 3-year for Sentinel-2.

Lastly, we extracted phenological metrics from smooth NDVI data. These metrics include the start of the growing season (SOGS) - the day of the year when the rising edge of the fitted curve reaches 20% of its seasonal amplitude, length of the growing season (LOGS) - the period between the start and the end of the growing season; base NDVI (B_NDVI) value – mean value of the lowest NDVI values to the left and right of the Peak of the Season, the peak of the growing season (POGS) - the day of the year when NDVI of the fitted curve reaches its maximum greenness; peak NDVI (P_NDVI) value – maximum value in the NDVI timeseries; and amplitude of NDVI (A_NDVI) - the difference between the maximum values and the base NDVI values per pixel (see S1 Fig for the graphical description of each phenological metrics) (32,55,60).

We further compared the phenological metrics across different land cover classes using a one-way ANOVA and the Kruskal-Wallis test (61). We stack the six extracted phenology metrics on top of the initial spectral and multispectral indices for their corresponding satellite imagery (see Table 2 for a summary of the metrics).

### 2.4 Training, Validation, and Test Datasets

Training data for land cover analysis involves selecting a sample of pixels from the image and using it to establish thresholds for delineating specific land covers on the ground (67). We extracted and stacked spectral, multispectral, and biophysical indices computed from the satellite imagery and the DEM, and then normalized them using Z-score normalization techniques (27). A supervised deep learning model typically requires a substantial amount of training data samples. However, it is often very time-consuming and costly to label the observed data and create training samples for each LULC class (38). Therefore, a series of training, test, and validation polygons were drawn across each class using Google Earth Imagery as the base map for the years 2010 and 2021, which were converted to raster based on the classes. The datasets for training, testing, and validation were generated by randomly selecting pixel patches from each corresponding train, test, and validation raster, with patches spaced at least 90 meters (i.e., 3 pixels) apart, ensuring class stratification across all satellite sensor types. This homogeneous patch sampling ensures that pixels are only derived from a region of consistent land cover. Specifically, for the co-available year (2021), the shape of each of the patches sampled and fitted into the model were as follows: (a) Landsat-8 full feature model is 7 (width) x 7 (height) x 26 (Bands), and 20 bands for the non-phenology models; (b) while Landsat-5 was trained with just the full feature model; and (c) Sentinel-2 full feature model is 21 (width) x 21 (height) x 26 (Bands), and 20 bands for the non-phenology models). This totaled up to 600 image patches per class, except for the Ghost Forests (310), for each satellite. The training, validation, and test data were split by a ratio of 7:1.5:1.5, stratified by class per image sensor, with the training set for the Landsat-5 model from 2010 imagery, while 2021 imagery for both Landsat-8 and Sentinel-2 models, except for the ghost forest classes, supplemented by 2019 samples, while 80% of the test and validation sets were from the 2019 imagery.

### 2.5 Classification method

CNN is a hierarchical feature detector, inspired by biological principles, with a classification capability based on contextual information (68). It is capable of learning highly abstract features and efficiently identifying objects through weight sharing and feature extraction (38,69–71). Due to this characteristic, CNN is well-suited for processing multiband remote-sensing image data, where pixels are arranged in a regular pattern (38). Fully trained full-feature CNN models were validated using the validation datasets and tested using the testing sets obtained from previous steps to classify forest classes and identify areas of decline.

The architecture includes four convolutional layers, each applying 32 to 64 filters (27, 72) with a kernel size of 3×3 (see Fig 3). An L2 regularization of 0.01 was used to prevent overfitting, given the large feature space relative to spatial context. The training was terminated at the epoch (each being a complete pass through the training data) corresponding to the lowest validation loss for each of the sensors (see S2 Fig), with a batch size of 32 batches (72) and a learning rate of 0.0001 based on the cross-validation performance. The filter, also known as the weight vectors, slides over the input vector (i.e., stacked image patches) to generate the feature map (73). This method, which involves sliding the filter horizontally and vertically, is known as a convolution operation (74). This operation extracts X features from the input image within a single layer, producing X filters and X associated feature maps (74). The output a_ij_ in the next layer for location (i,j) is computed after applying the convolution operation using the formula given by (71) as shown below:

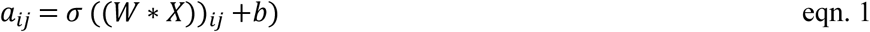

where X is the input given to the layer, W is the filter or kernel that moves over the input, b is the bias, * signifies the convolution operation, and σ indicates the non-linearity added in the network. The layer that carries out the classification was converted to a convolutional layer. The convolution kernel is chosen so that its dimensions match those of the input layer (71).

**Figure 3.**
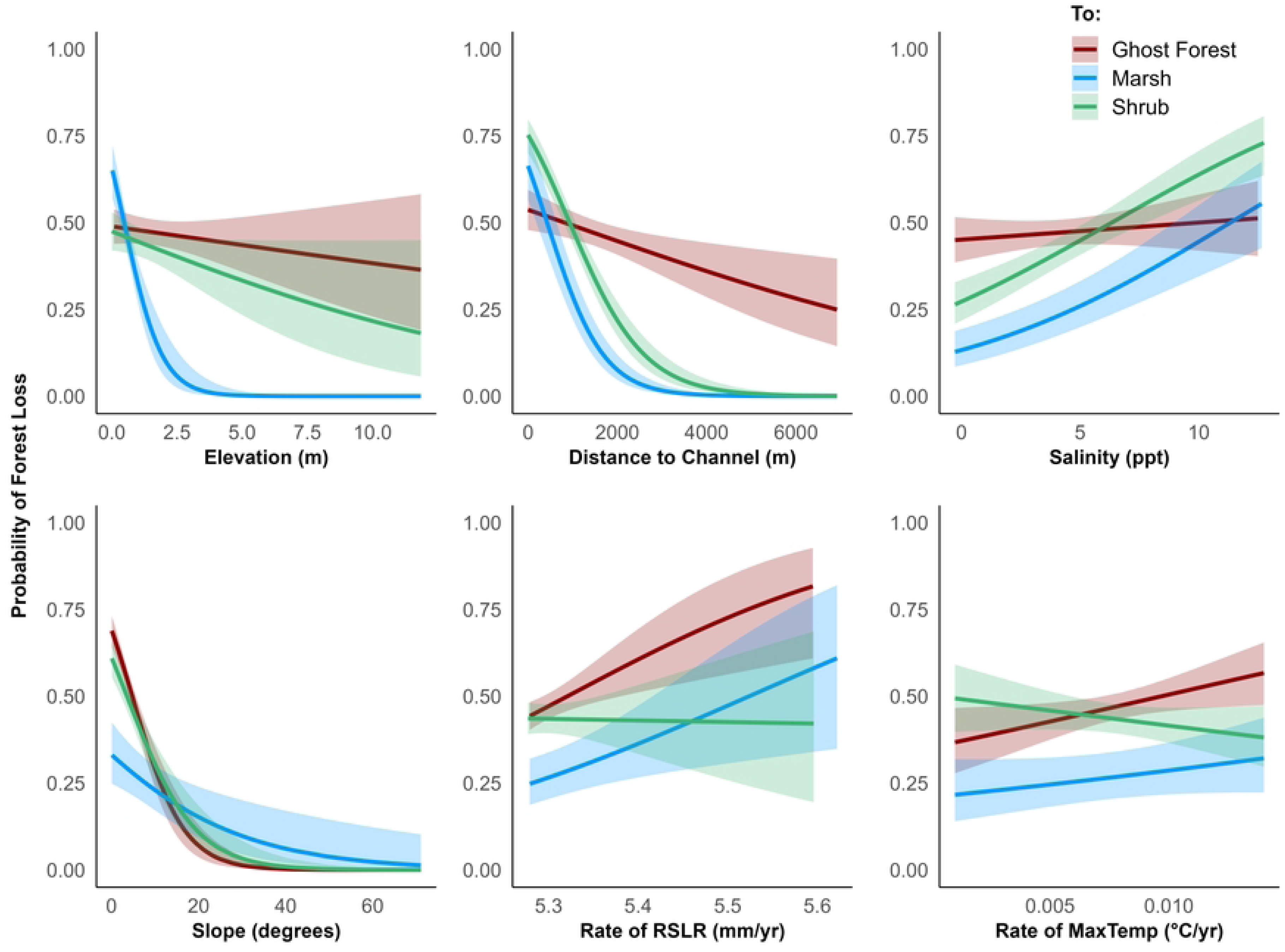
The architecture of the Convolutional Neural Network (CNN) used for image classification consists of multiple convolutional layers followed by a fully connected output layer for final prediction.

To evaluate the classification results across the various satellite imagery (given the difference in temporal, spectral, and spatial resolution), we computed the confusion matrix table, which was used to calculate the Accuracy (eqn. 2), User Accuracy (eqn. 3), Producer Accuracy (eqn. 4), and F1 score (eqn. 5)(27, 32) as follows:

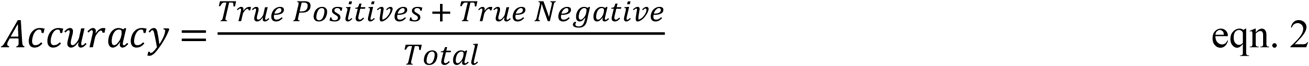

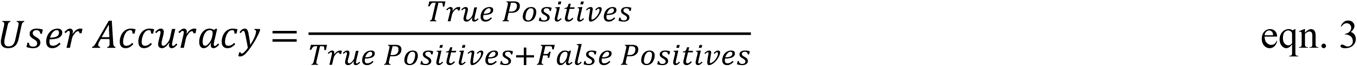

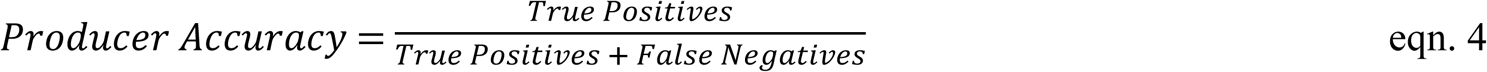

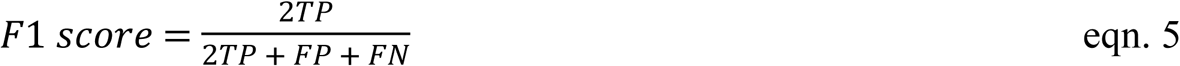

A confusion matrix compares predicted and observed class labels, and it summarizes model outcomes into four categories: true positives (TP), true negatives (TN), false positives (FP), and false negatives (FN). The user’s accuracy (Precision) is the proportion of correct predictions for each predicted class in the test data. In contrast, the producer’s accuracy (Recall) represents the proportion of correct predictions for each actual class (75), while the F1 score is the harmonic mean of both.

An image patch to be classified is fed into the input layer, and the output is the predicted land cover class label for each pixel, computed using the extracted features from the image. Furthermore, we utilized the permutation feature importance (PFI) to evaluate the significance of each feature in the model. PFI measures the degree to which a trained model relies on each feature (76). After training, the values of a single feature are randomly shuffled, and a drop in model performance indicates the importance of that feature. If the error (Mean Absolute Error (MAE), i.e., the average magnitude of errors between predicted values and actual (observed) values) increases significantly when a feature is shuffled, it indicates that the feature is important. Since shuffling the features leads to an increase in the error metric, it means that these features are contributing positively to the model’s performance (77). The size of the values (how far above zero they are) indicates the level of importance. Larger MAE values indicate that the feature plays a more significant role in the model’s predictions.

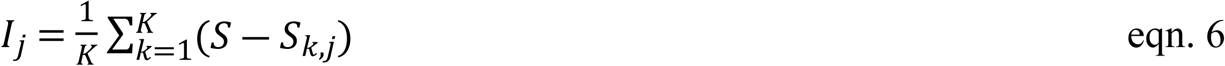

Where I is used to index the features, j represents the total number of features, 1_j_ is the importance of the j-th feature, S is the baseline performance of the model (i.e., accuracy), is the score after permuting the j-th feature in the k-th repetition, and K is the number of permutations, which in this case is 20. We calculate the average MAE over the total 20 runs and display the variation as a standard deviation.

We also conducted post-processing of our final classified map to remove obvious classification errors. Additionally, to facilitate cross-product comparisons of coastal forests, we compared our results with the National Land Cover Database (NLCD) for 1985 and 2021. NLCD uses a complex system comprising approximately 20 land-cover classes (78). For comparison, we consolidated NLCD pixels labeled evergreen forest, deciduous forest, shrub/scrub, mixed forest, and woody wetland as forests. We considered the NLCD emergent herbaceous wetlands to be marshes (32). In our study, we also merged the forest, ghost forest, and shrub classes into a single class to represent forests for comparison.

### 2.6 Drivers of Forest Retreats

We examined coastal forest loss between 1985 and 2021 by identifying pixels classified as forest in 1985 that have since transitioned to other land cover types, such as shrub, wetland, or ghost forest. This is done only for the Landsat Imagery because it’s long-term, while Sentinel-2 was used only for the 2021 image comparison with Landsat 8. Using the slope and elevation computed from the CONED DEM (66), we assessed the topographical impact on forest loss and the formation of ghost forests. We also tested the relationship between distance to channel, salinity, climatic variables, and forest loss. We examined three major pathways of forest change (i.e., forest-to-ghost forest, forest-to-shrub, and forest-to-marsh). For each pathway, we used logistic regression to assess the factors driving the dynamic patterns of coastal forest change (see Table 3). A total of 1,000 points were randomly generated across all pixels/areas, stratified by change types (i.e., binary change), using a 500-meter spatial exclusion buffer to reduce autocorrelation as employed by Ury et al. (2021) (9). Other environmental predictors, such as salinity, distance to channel, and climate variables, were also employed. A k-fold cross-validation approach was employed to fit a binomial logistic regression model in which binary coastal forest change is modeled as a function of the predictors, including distance to water, salinity, relative sea level rise, and elevation (Table 3).

**Table 3.**
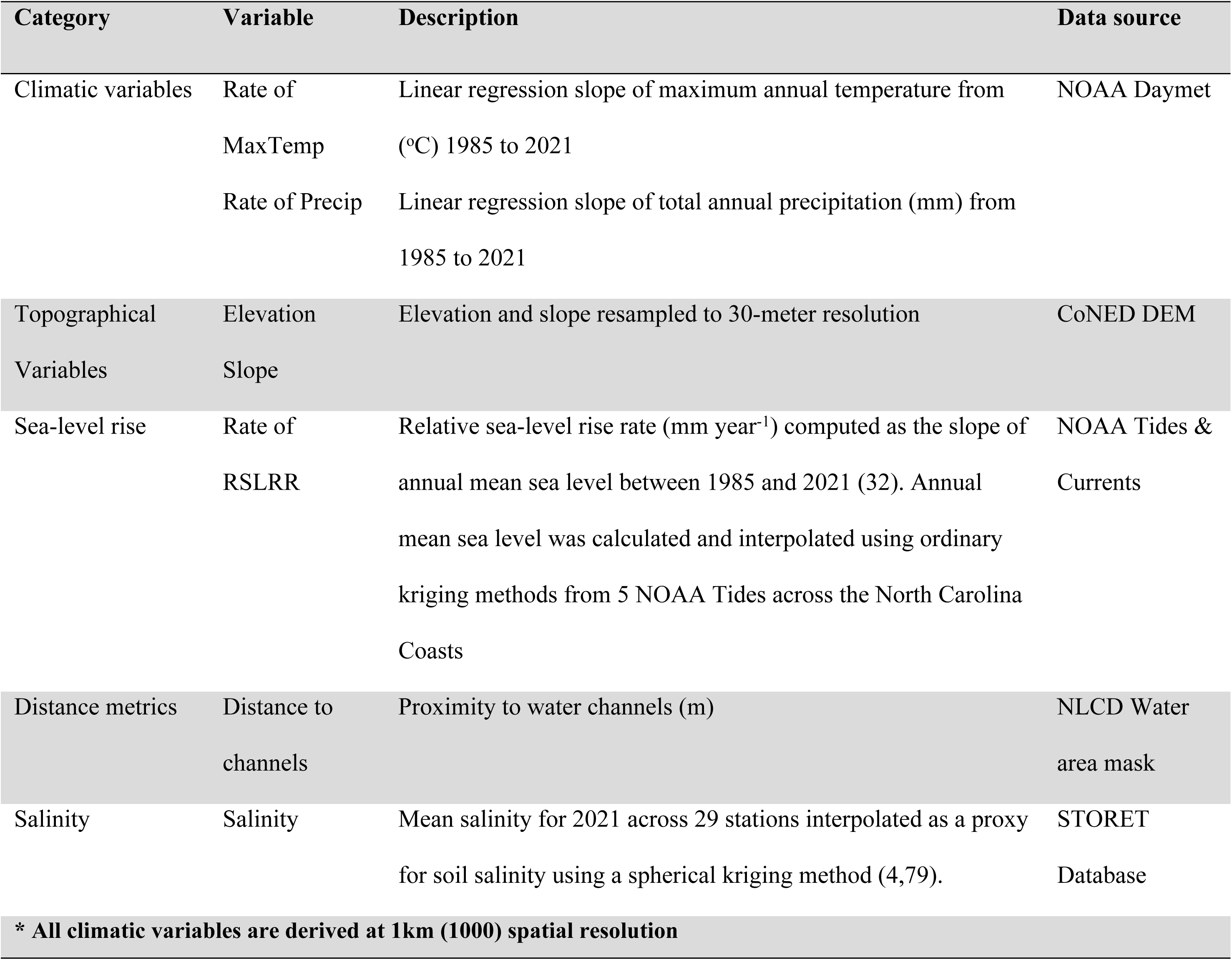
Potential predictors of forest change.

| Category | Variable | Description | Data source |
| --- | --- | --- | --- |
| Climatic variables | Rate of | Linear regression slope of maximum annual temperature from (°C) 1985 to 2021 | NOAA Daymet |
|  | MaxTemp |  |  |
|  | Rate of Precip | Linear regression slope of total annual precipitation (mm) from 1985 to 2021 |  |
| Topographical Variables | Elevation | Elevation and slope resampled to 30-meter resolution | CoNED DEM |
|  | Slope |  |  |
| Sea-level rise | Rate of | Relative sea-level rise rate (mm year<sup>-1</sup>) computed as the slope of annual mean sea level between 1985 and 2021 (32). Annual mean sea level was calculated and interpolated using ordinary kriging methods from 5 NOAA Tides across the North Carolina Coasts | NOAA Tides & |
|  | RSLRR |  | Currents |
| Distance metrics | Distance to channels | Proximity to water channels (m) | NLCD Water area mask |
| Salinity | Salinity | Mean salinity for 2021 across 29 stations interpolated as a proxy for soil salinity using a spherical kriging method (4,79). | STORET Database |

**\* All climatic variables are derived at 1km (1000) spatial resolution**

## 3 Results

### 3.1 Comparison of phenological metrics between landcover classes based on the preprocessing steps

Coastal land cover types exhibit contrasting patterns of land surface phenology (Fig 4). For example, as obtained from the training data, the Peak NDVI value for both Landsat and Sentinel images reaches 0.88 ± 0.02 in the forest, which is higher than that of marsh (0.57 ± 0.08), shrub (0.72 ± 0.04), and ghost forest (0.83 ± 0.02) vegetation. Another thing that stands out in Figure 4 is the annual variability of the forest class (D) for the Landsat imagery, which differs from that of the other classes and may be related to differences in reflectance values between Landsat 5 and Landsat 8.

**Figure 4.**
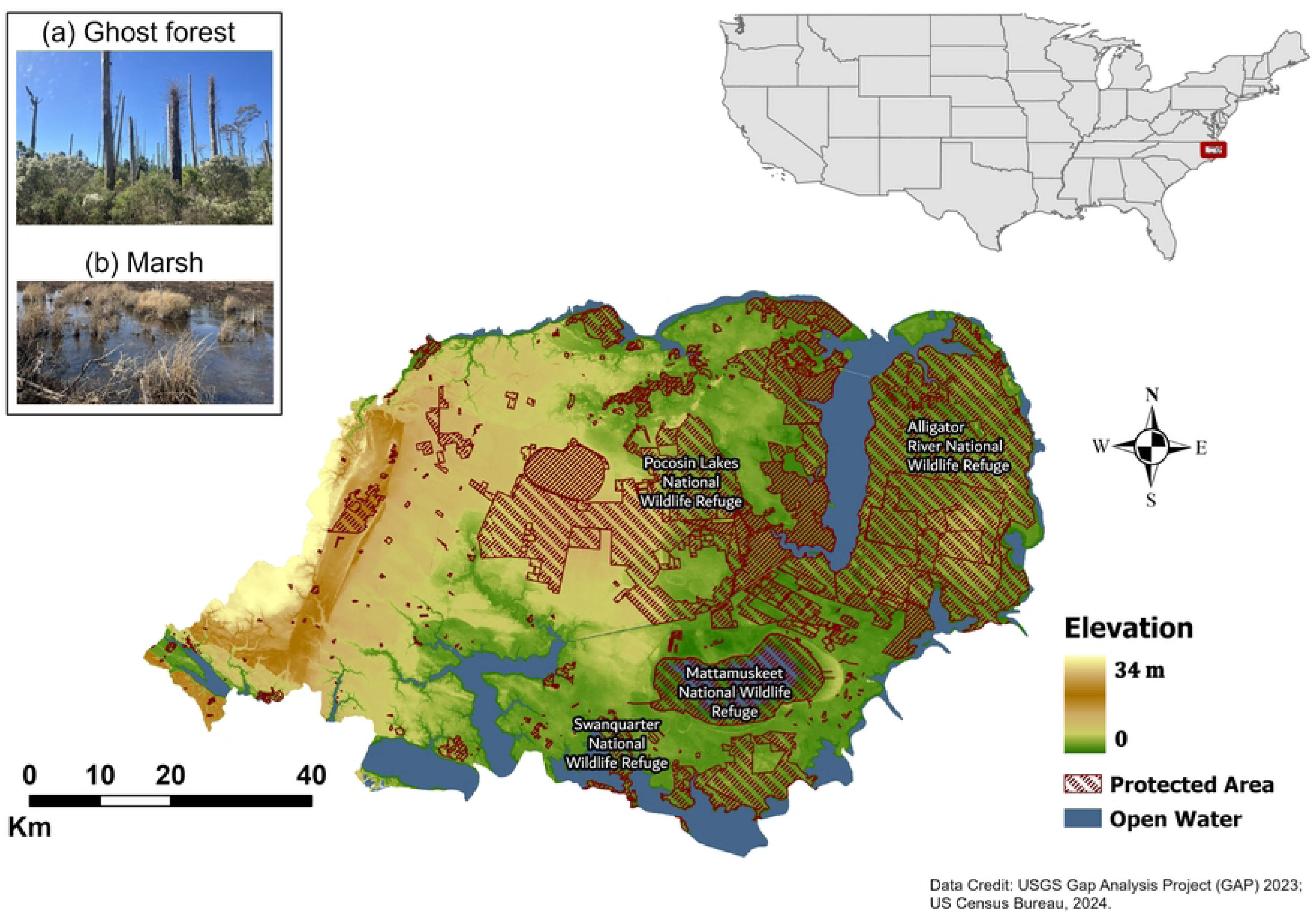
Annual vegetation patterns estimated from training data for marsh (A), shrub (B), ghost forest (C), and forest (D) were estimated using Landsat NDVI time series, while E-H represent the Sentinel 2 NDVI trend. Plots show the mean DOY NDVI ± SD data at three time steps.

Additionally, the Landsat Forest (103 ± 27 DOY) has the earliest start of growing season (SOGS) value, and the shrub (123 ± 23 DOY) has the latest SOGS value, while the other classes are in between. We also observed a distinct difference between the satellites’ SOGS values for ghost forests, with the start of the growing season for the Sentinel-2 images occurring 17 days earlier than the Landsat images, and the forest starting 2 days earlier. Differences are evaluated using a one-way ANOVA, and the results indicate that all phenology metrics differ among the four vegetation classes for the Sentinel-2 and Landsat 8 images (see Table 4). The ANOVA results show that the interclass difference at the start of the growing season is significant (p ≤ 0.001), as well as for all other metrics. However, all other metrics have a low R^2^ (R^2^ ≤ 0.11, i.e. while the difference between classes is statistically significant, only 10-21% of the variability in SOGS, POGS, A_NDVI can be explained by class) except for the Base NDVI (R^2^ = 0.79) and Peak NDVI (R^2^ = 0.86) for Landsat 8 images indicating that a large proportion of the variation these metrics can be explained by the class difference. A similar pattern was observed in the extracted metrics from the Sentinel-2 images, but a lower R-squared value (R^2^ ≤ 0.019) in SOGS, POGS, and LOGS with A_NDVI (R^2^ = 0.28), Base NDVI (R^2^ = 0.59), and Peak NDVI (R^2^ = 0.83). There is also a variation in the length of the growing season for all land cover types except ghost forests (approximately 225 days) between the two satellites (see Table 4). However, the base NDVI, Amplitude of NDVI, and Peak NDVI metrics for both satellite sensors are similar.

**Table 4.**
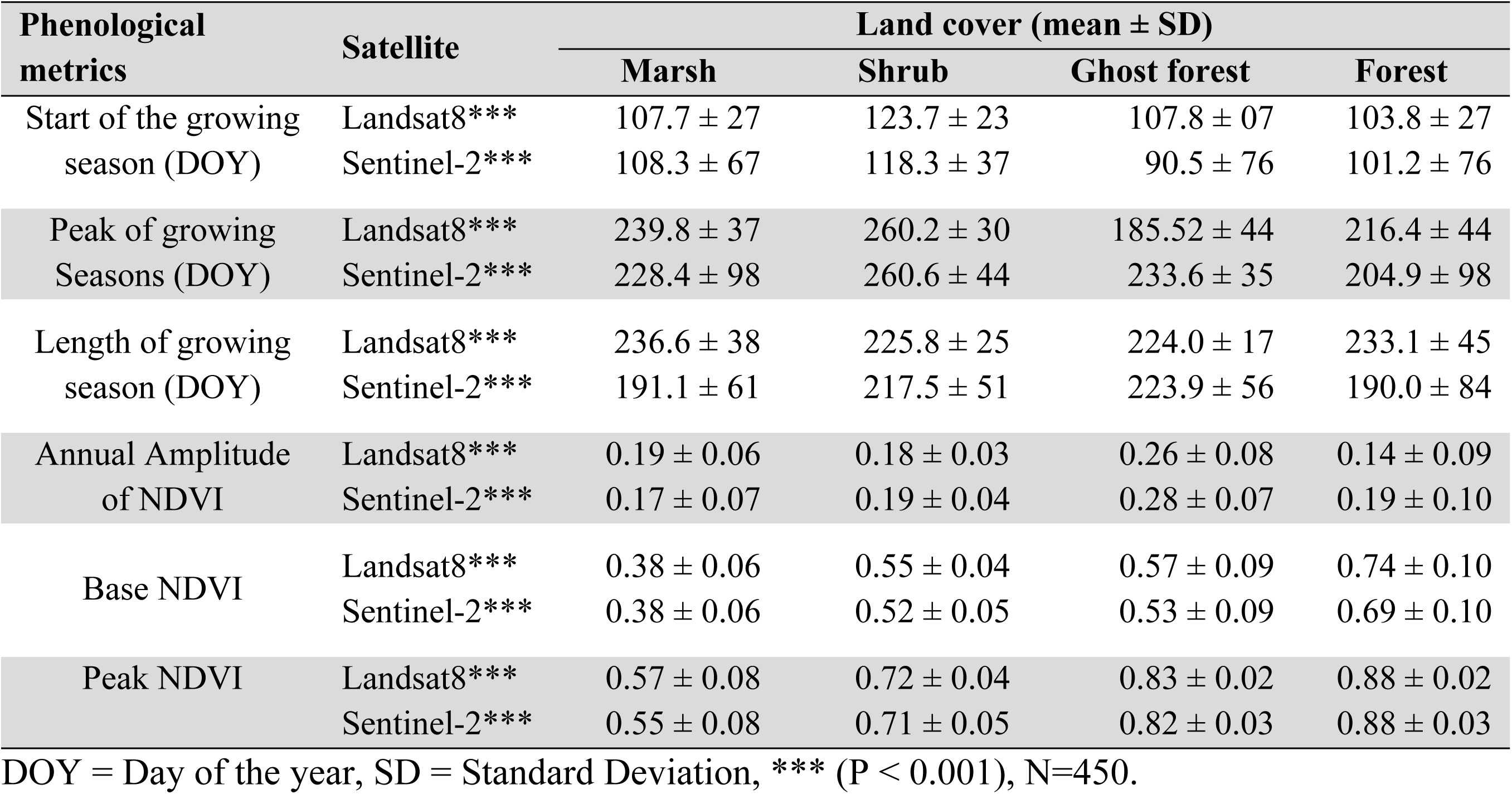
Phenological results for four classes estimated from the training samples.

| Phenological metrics | Satellite | Land cover (mean ± SD) | | | |
| --- | --- | --- | --- | --- | --- |
|  |  | Marsh | Shrub | Ghost forest | Forest |
| Start of the growing season (DOY) | Landsat8*** | 107.7 ± 27 | 123.7 ± 23 | 107.8 ± 07 | 103.8 ± 27 |
| | Sentinel-2*** | 108.3 ± 67 | 118.3 ± 37 | 90.5 ± 76 | 101.2 ± 76 |
| Peak of growing Seasons (DOY) | Landsat8*** | 239.8 ± 37 | 260.2 ± 30 | 185.52 ± 44 | 216.4 ± 44 |
| | Sentinel-2*** | 228.4 ± 98 | 260.6 ± 44 | 233.6 ± 35 | 204.9 ± 98 |
| Length of growing season (DOY) | Landsat8*** | 236.6 ± 38 | 225.8 ± 25 | 224.0 ± 17 | 233.1 ± 45 |
| | Sentinel-2*** | 191.1 ± 61 | 217.5 ± 51 | 223.9 ± 56 | 190.0 ± 84 |
| Annual Amplitude of NDVI | Landsat8*** | 0.19 ± 0.06 | 0.18 ± 0.03 | 0.26 ± 0.08 | 0.14 ± 0.09 |
| | Sentinel-2*** | 0.17 ± 0.07 | 0.19 ± 0.04 | 0.28 ± 0.07 | 0.19 ± 0.10 |
| Base NDVI | Landsat8*** | 0.38 ± 0.06 | 0.55 ± 0.04 | 0.57 ± 0.09 | 0.74 ± 0.10 |
| | Sentinel-2*** | 0.38 ± 0.06 | 0.52 ± 0.05 | 0.53 ± 0.09 | 0.69 ± 0.10 |
| Peak NDVI | Landsat8*** | 0.57 ± 0.08 | 0.72 ± 0.04 | 0.83 ± 0.02 | 0.88 ± 0.02 |
| | Sentinel-2*** | 0.55 ± 0.08 | 0.71 ± 0.05 | 0.82 ± 0.03 | 0.88 ± 0.03 |

DOY = Day of the year, SD = Standard Deviation, \*\*\* ( $P < 0.001$ ), N=450.

### 3.2 Classification Results and Evaluation

We assessed the model’s accuracy using the user’s, producer’s, and F1 score metrics, as well as the loss curve. Firstly, as shown in S2 Fig, the training and validation accuracies increased rapidly during the initial epochs and plateaued at around 0.94, indicating that the model effectively learned the underlying patterns for both Sentinel 2 and Landsat. Correspondingly, the training and validation loss decreased steadily, stabilizing at approximately 0.4 by the final epoch. The close alignment between the training and validation curves for both Sentinel-2 and Landsat-8, with minimal divergence, suggests that the model achieved high predictive performance and good generalization on unseen datasets without significant overfitting.

For the Landsat dataset, the classification results for all land cover types demonstrated a high classification accuracy (weighted average accuracy = 94% & F1 Score = 93%), indicating that most instances of each land cover class were correctly classified into their true class with few misclassifications. For the Ghost Forest class, the results show the presence of false negatives, instances where areas that should have been classified as Ghost Forest were instead misclassified as forest (11%) and Shrub (5.5 %), suggesting underestimation and potential overestimation by the forest samples, with about 5.1% false positives in the model estimation. Additionally, 8% of the marsh class was misclassified as water.

Similar to Landsat, the producer’s accuracy for Sentinel imagery ranged from 90% for ghost forest to 99% for water, emphasizing the model’s overall consistency. Sentinel achieved an overall accuracy of 97%, indicating a high level of agreement (see Table 5). However, 11% of ghost forest pixels were misclassified as shrub, while 7% of water pixels were misclassified as marsh. There is an 8% false positive for the ghost forest class, indicating an overestimation of ghost forest areas, i.e., non-ghost forest areas (forests, shrubs, and cultivated) were classified as such

**Table 5.**
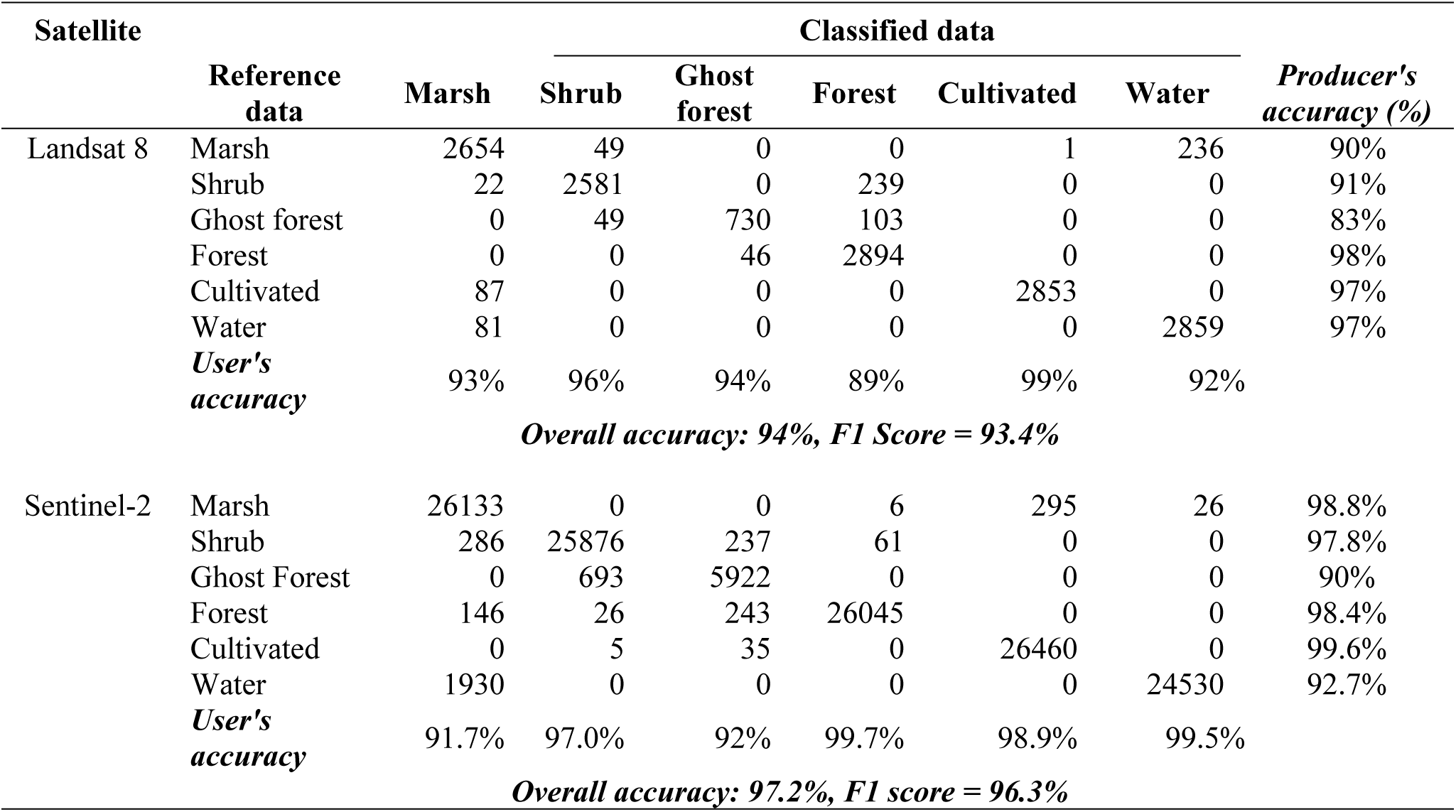
The confusion matrix showing the classification accuracy of land-cover maps produced by the Landsat 8 and Sentinel 2 satellites in 2021 using all 26 features.

|  | Reference data | Marsh | Shrub | Ghost forest | Forest | Cultivated | Water | Producer's accuracy (%) |
| --- | --- | --- | --- | --- | --- | --- | --- | --- |
| Landsat 8 | Marsh | 2654 | 49 | 0 | 0 | 1 | 236 | 90% |
|  | Shrub | 22 | 2581 | 0 | 239 | 0 | 0 | 91% |
|  | Ghost forest | 0 | 49 | 730 | 103 | 0 | 0 | 83% |
|  | Forest | 0 | 0 | 46 | 2894 | 0 | 0 | 98% |
|  | Cultivated | 87 | 0 | 0 | 0 | 2853 | 0 | 97% |
|  | Water | 81 | 0 | 0 | 0 | 0 | 2859 | 97% |
|  | **User's accuracy** | 93% | 96% | 94% | 89% | 99% | 92% |  |
| **Overall accuracy: 94%, F1 Score = 93.4%** |  |  |  |  |  |  |  |  |
| Sentinel-2 | Marsh | 26133 | 0 | 0 | 6 | 295 | 26 | 98.8% |
|  | Shrub | 286 | 25876 | 237 | 61 | 0 | 0 | 97.8% |
|  | Ghost Forest | 0 | 693 | 5922 | 0 | 0 | 0 | 90% |
|  | Forest | 146 | 26 | 243 | 26045 | 0 | 0 | 98.4% |
|  | Cultivated | 0 | 5 | 35 | 0 | 26460 | 0 | 99.6% |
|  | Water | 1930 | 0 | 0 | 0 | 0 | 24530 | 92.7% |
|  | **User's accuracy** | 91.7% | 97.0% | 92% | 99.7% | 98.9% | 99.5% |  |
| **Overall accuracy: 97.2%, F1 score = 96.3%** |  |  |  |  |  |  |  |  |

The Landsat 5 Classifier is trained using 2010 and 2011 images, achieving an overall accuracy of 94% with an F1 score of 93.8%. Landsat 5 has the lowest accuracy for users and producers in the Ghost Forest class compared to Landsat 8 and Sentinel-2, which could be a result of a very small training dataset for the Ghost Forest class.

Here, the result indicates that the phenological metrics have the highest importance on the model prediction, especially the Base NDVI (B_NDVI), amplitude of NDVI (A_NDVI), and peak growing season (POGS) for the Landsat 8 image, followed by the winter spectral indices, with the NIR band leading to an approximately 0.10 mean absolute error in classification accuracy when shuffled (Fig 5). On a per-class level, the results show that multispectral indices (mNDWI) aid in distinguishing water from every other class, while a combination of Amplitude of NDVI, mNDWI, winter red, and blue helps in separating marsh and every other class, as indicated by the model fit error. For the Ghost Forest class, winter blue and peak NDVI help in separating from other vegetation classes (see S3 Fig).

**Figure 5.**
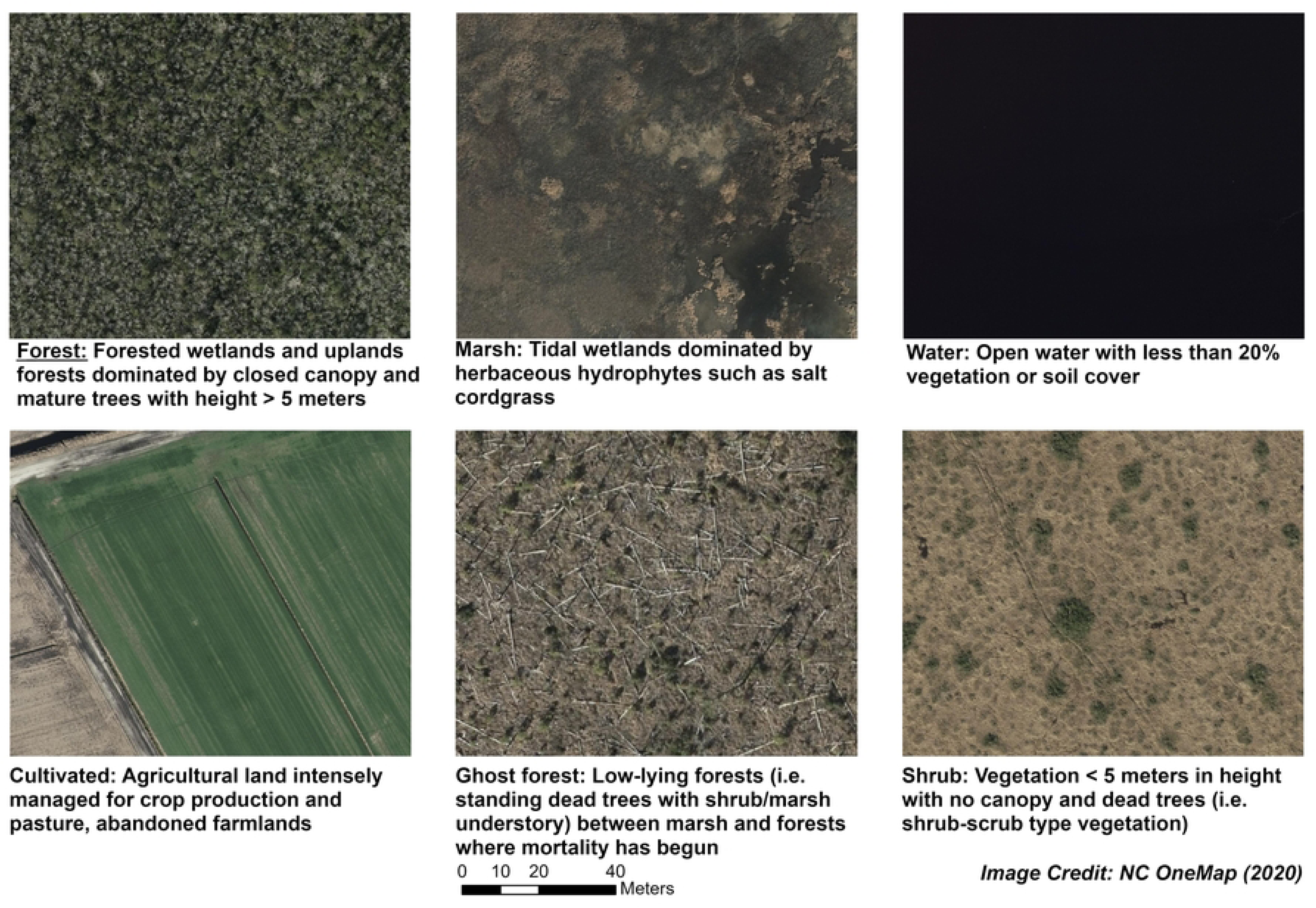
(a) User’s Accuracy and (b) Producer’s accuracy of classification results from Landsat 5, Landsat 8, and Sentinel 2; (c) Landsat 8’s and (d) Sentinel-2’s permutation feature importance value - the red and blue colors represent the spectral indices for winter and summer, respectively while green represents multispectral indices gray for topographical and orange represents the phenological indices.

In contrast, in the Sentinel-2 full feature prediction, multispectral indices have the most significant impact on the prediction, specifically the modified Normalized Difference Water Index (mNDWI), resulting in a 0.17 reduction in accuracy when shuffled. The Peak growing season (POGS), Amplitude, and Base NDVI metrics also contribute significantly to the model prediction, with an average of 0.085 increase in error when removed from the model. On a per-class level, mNDWI distinctly aids in the separation of water and marsh from every other class, while wintergreen, short-wave infrared, and peak NDVI aid in separating ghost forests and other vegetation classes. TCP greenness, the start of the growing season, and Base NDVI have the highest separation power for the forest class (see S4 Fig). Lastly, for both satellites, phenology is the major feature that aids in the separability of shrub from other classes (see S3-S4 Figs). The non-phenology model (i.e., excluding phenology) shows that without phenology, Sentinel-2 held only a marginal advantage over Landsat 8 (test accuracy 0.967 vs. 0.948; gap = 0.019), while Sentinel-2’s advantage increases over Landsat 2 (0.972 vs. 0.941; gap = 0.031). Sentinel-2’s accuracy improved with the addition of phenology, while Landsat 8’s accuracy declined (McNemar’s test, p < 0.001 for both sensors).

### 3.4 Forest Change Analysis

Given the limited time series data available from Sentinel-2, the forest change analysis primarily focused on Landsat Imagery. In the study area of 624,892 ha, 48.8% (304,729 ha) was forest in 1985, which reduced to 43.2 % (269,970 ha) in 2021 (see Fig 6c). Over 36 years, the Albemarle Peninsula study area, which includes private land as well as Pocosin Lakes National Wildlife Refuge and other preserved areas, experienced a net loss of 21.1 % in forest cover, or approximately 64,220 ha. Approximately 10% (30,336 ha) were converted to cultivated land. Additionally, 33,884 ha (53%), i.e., 940 ha yr^-1^ transitioned to either shrub (13423 ha, 372 ha yr^-1^), marsh (9690 ha, 269 ha yr^-1^), water (1364 ha, 38 ha yr^-1^), and ghost forest (9405 ha, 261 ha yr^-1^) by 2021. Specifically, 23,876 ha of forest were lost between 2010 and 2021 to marsh, ghost forest, and shrub, which is 1.5 times higher than the 16,968 ha loss between 1985 and 2010.

**Figure 6.**
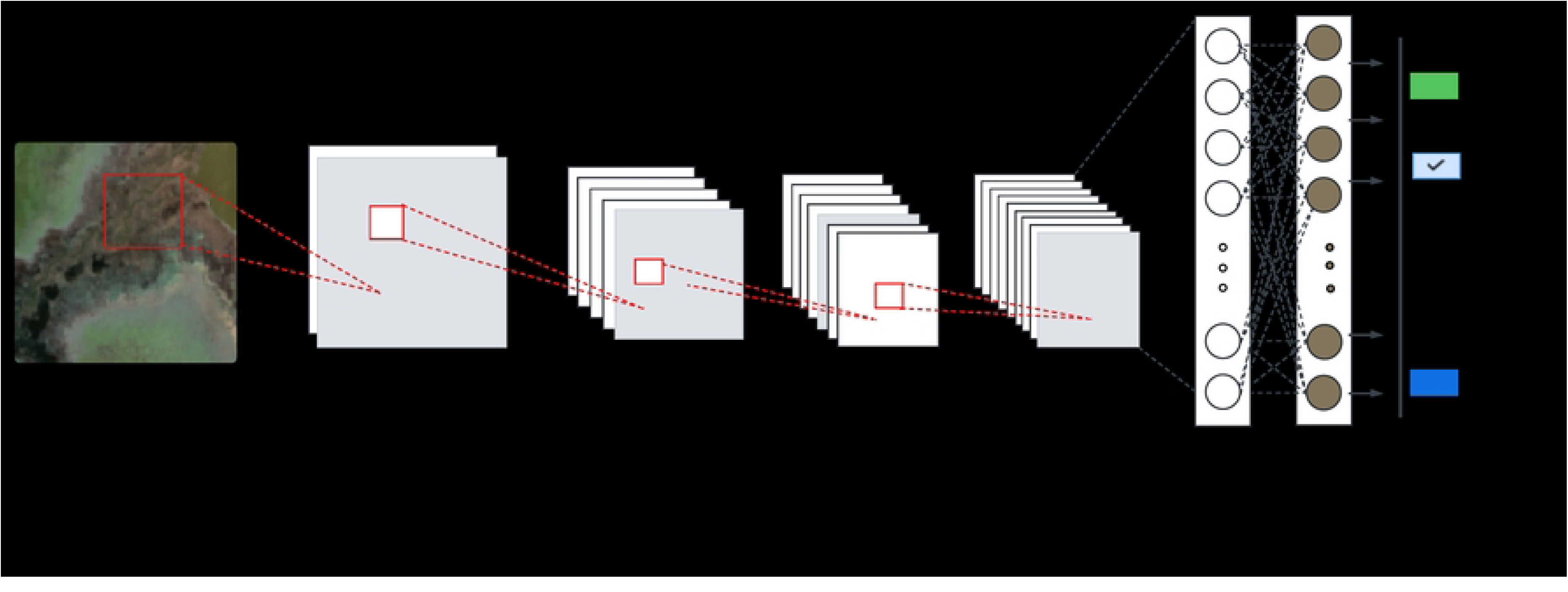
(a) 1985 Coastal land cover, (b) 2021 land cover map, (c) pattern of coastal land cover change, (d-i) Comparison of classified data to existing land cover products. The maps presented in g-i plots are our maps generated in 1985 and 2021, followed by the differenced map (i.e. area of forest-marsh gain or loss). The corresponding NLCD products are shown in d-f, followed by the differenced map, which shows the area of marsh-forest gain and loss from 1985 to 2021.

Overall, we found land cover change was dynamic across our study area between 1985 and 2021 (Fig. 6). We quantified a 28% increase in marsh between 1985 (26228 ha) and 2021 (33611 ha). Forests accounted for 43% of the marsh area increase between 2010 and 2021, and 60% of the marsh increase between 1985 and 2021. Much of the remaining increase in marsh was shrubland in 1985. Ghost forests had a net gain of 7561 ha (687 ha yr^-1^) between 2010 and 2021, which is 2.5 times higher than the estimated net gain in ghost forest area between 1985 and 2010 (3087 ha). Our results show that forest-shrub transition (i.e., shrubland gain) was higher relative to forest-marsh and forest-ghost forest loss between 1985 and 2021, with a net gain of 24% (3638 ha) between 1985 and 2010 and a net gain of 10% (1877 ha) between 2010 and 2021. Lastly, we found some areas of forest gain, and cultivated land accounted for 47% of reforestation, mostly abandoned farmlands, followed by shrubland (21%) between 1985-2021.

We compared our results to existing products, such as the NLCD data. As presented in Fig 6, there is a close overlap in the forest-wetland boundary area loss. However, our study shows approximately 71% more area of forest-wetland loss compared to the NLCD estimated loss.

### 3.5 Patterns and Predictors of Forest Change

In our analysis, each coefficient (estimate) represents the change in log odds of the outcome for a 1-unit increase in the predictor, holding other variables constant. Distance to the channel was particularly predictive of transitions between forest and shrub, and between forest and marsh. The result of the logistic regression model showed that for each additional meter from the channel, the odds of forest loss to marsh decreased by approximately 0.16% (*β* = -0.0016, *p* < .0001) and the odds of forest loss to shrubs decreased by 0.13% (*β* = -0.0013, *p* < .0001), see Fig. 7. Salinity was one of the most significant predictors of ghost forest formation as well as forest loss to marsh. With a significant coefficient (p < .01), the model showed that every unit increase in salinity leads to a 0.0628 unit increase in the log odds of forest loss to ghost forest (β = 0.0628, p < .05) and even more loss to shrub (β = 0.1637, p < .0001). Like the distance to the channel, a 1-meter increase in elevation leads to a 1.54 unit decrease in the log odds of forest-to-marsh change (β = -1.536, p < 0.0001). Although slightly weaker statistically, the log odds of forest transition increased by 9% for ghost forests (β = 0.0906, p < 0.05) and 16% for shrubs (β = 0.1654, p < 0.05) per unit decrease in elevation, respectively. Ghost forest formation was significantly and positively influenced by RSLRR (β = 8.120, p < .0001) as was forest-marsh loss (β = 7.69, p < .0001). An increasing trend in the annual maximum temperature (slope) was observed, with the slope of annual maximum temperature strongly and positively influencing forest loss to ghost forest (β = 72, p < .0001), indicating that the warming trend leads to higher forest loss. However, it did not have any significant effects on the observed shrub and marsh transgression.

**Figure 7.**
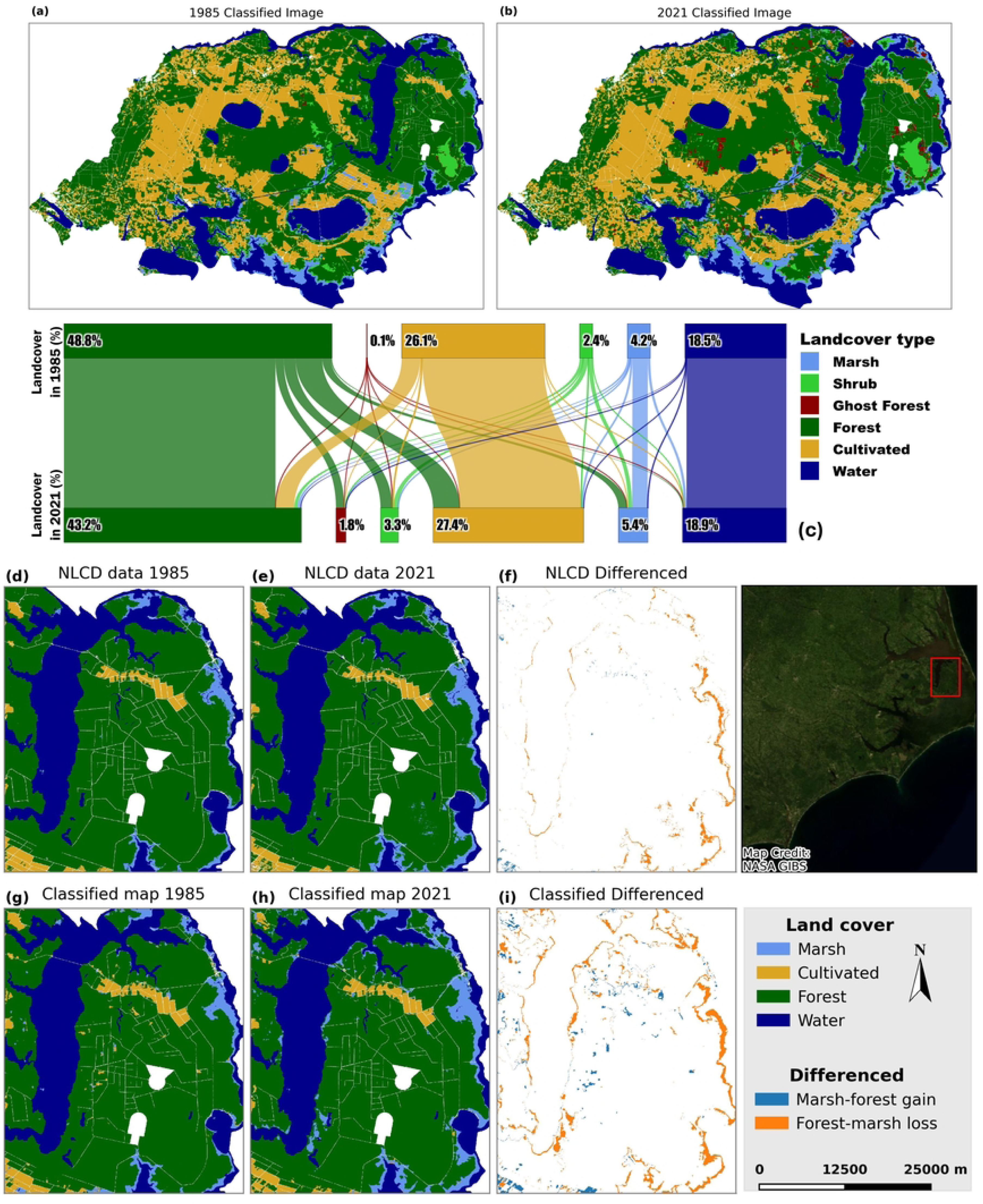
Predicted probability of forest loss for each driver. The lines represent model-predicted probabilities of each change type across all environmental drivers with 95% confidence intervals (shaded areas).

## 4 Discussion

### 4.1 Phenological Insights from Coastal Vegetation Mapping with Landsat and Sentinel2

With accelerating rates of forest loss resulting in expansive ghost forests and shrub zones, our study demonstrates how remote sensing combined with CNN models can improve separability between vegetation classes and map land cover with spatial precision. Our findings support the use of DL models and phenology to improve land cover mapping and change analysis using Landsat and Sentinel imagery. We observed that both sensors demonstrated high classification performance across all classes. However, Sentinel-2 (F1 score = 91%) showed slightly higher accuracy for the ghost forest class compared to Landsat-8 (F1 score = 88%), possibly due to its higher resolution, distinct phenological attributes, and more training samples.

Our results demonstrate that combining winter and summer composite images with phenological and multispectral indices significantly improves the classifier’s ability to distinguish among the shrub, forest, and ghost forest categories. This improvement is due to seasonal differences in reflectance, which are better captured when images from both seasons are included, rather than using only summer images (see Fig 5 and S3 Fig). Accordingly, our approach to accounting for the phenological differences of coastal vegetation enables improved mapping accuracy, with phenological metrics contributing 36% (Fig 5) to the Sentinel 2 model, following the contribution of multispectral indices (37%) and 40% to the Landsat 8 model accuracy (Fig. 5), followed by Winter indices at 29%. Earlier research indicated that incorporating phenology yields higher accuracy than traditional biseasonal methods, suggesting that land surface phenology is crucial for constraining intra-class variations in complex landscapes (27,32,33,45) and for separability between spectrally similar tree species (39). In fact, phenology was the highest-performing predictor across all classes for both sensors (Fig 5), especially for the Shrub class. However, winter red and green also contributed to ghost forest class separation for Landsat 8 (S3 Fig) and Sentinel-2 (S4 Fig), respectively. As reported by Ury et al. (2021) (9), distinguishing between shrub and ghost forest was more challenging for their classifier due to spectral similarities, which further highlights the importance of phenology and the CNN used in our study. This result further shows the strong overlap between ghost forest, shrub, and forest as revealed in the confusion matrix (Table 5). Additionally, despite the heavy reliance on phenology, Landsat 8’s overall test accuracy is slightly lower with phenology included than without it, which further could mean that Landsat 8’s 16-day revisit interval may carry higher estimation variance that is sparser and noisier compared to Sentinel-2’s 5-day revisit period, which produces phenology metrics with distinguishable signals. However, Landsat compensates heavily for missing phenology in the non-phenology model by relying on correlated signals remaining in the spectral and multispectral indices (see S5(a)).

Our results underscore that Sentinel-2 data enables better mapping of forests, especially the ghost forest class, as revealed in the classification accuracy (Fig 5). This offers an opportunity for deep learning-based harmonization of Landsat and Sentinel-2 (80) or for using the output features from Sentinel-2 models to fine-tune or pre-train models for Landsat 8, potentially improving its classification accuracy for ghost forest or other vegetation transition mapping in future studies. Taking a closer look at phenology, we observed high variability in the phenological metrics for each vegetation class across the two satellite sensors (Table 4), which could be attributed to the mix of deciduous and evergreen trees. For instance, the forest class includes some planted pine plantations within the study site, alongside deciduous and evergreen forests that exhibit varying phenology throughout the year. Additionally, a few valid observations were available when clouds were masked out, and these were slightly improved by the 3-year composite.

Furthermore, the total land area classified for each class varies slightly between the two sensors, which may be due to the mixed-pixel effect or differences in band placement. For instance, the water area estimated by Sentinel-2 is about 5.5% lower than Landsat estimates, which may be due to a mix of water and non-water features in some pixels, while Sentinel-2 captures finer details, thereby reducing overestimation.

Comparing our results to the work of Chen and Kirwan (2022) (32), who have initially employed phenology for coastal vegetation mapping, our overall accuracy is higher, but their producers’ accuracy value for the Ghost Forest class (producer’s accuracy = 90%, user’s accuracy = 91.2%) is slightly higher than what we observed in our analysis (producer’s accuracy = 83%, user’s accuracy = 94%). A possible explanation for the difference, aside from not using the same training dataset, is that they do not have a Shrub class. In our study and previous studies (9), the ghost forest class was often misclassified as shrub, likely due to similarities in spectral signatures between dead tree stands and early successional vegetation. Additionally, about 11% of ghost forest pixels were misclassified as forests, as also observed by previous studies (4,32).

### 4.2 Comparing existing products and previous studies

While our study employed both Landsat and Sentinel-2, previous studies around the APP had only used Landsat imagery, which will serve as our basis of comparison (Table 6). The survey conducted by Ury et al. (2021) (9) using a traditional classifier achieved an overall accuracy of 92.2% for the Landsat 8 classifier, which is lower than our classifier at 94.3%. Smart et al. (2020) (4), on the other hand, show a CV-kappa (cross-validated Cohen’s kappa (CV-kappa) measures agreement between predicted and observed classifications while accounting for chance agreement) value of 0.60, which is lower than our kappa value of 0.95. While Smart et al. (2020) (4) also employed the traditional classifier (random forest model), another possible explanation could also be that their images were from 2014, while ours used 2021 images, indicating a 7-year gap.

**Table 6.** Comparing classification results and changed areas to existing studies along the Albemarle Peninsula and existing products.

| Product/Studies | Criteria (ha/Yr) |  |  |
| --- | --- | --- | --- |
|  | Forest loss<br>(Forest-Marsh) | Forest loss<br>(Forest-Ghost Forest) | Forest gain<br>(Shrub-Forest) |
| NLCD (81) | 166 | --- | --- |
| Ury et al. (2021) (9) | 259 | 328 | 266 |
| Martinez et al. (2024)<br>(62) | 145 | 188 | --- |
| Study result | 269 | 261 | 109 |
| *Note: The Ury et al. (2021) study includes the spatial extent of all pixels classified as forest-ghost loss or forest-marsh loss from 1985 to 2019. Our forest-marsh class is equivalent to the Martinez et al. (2021) 's Abrupt Transition (scenario 1: ecosystem is unstable and transitioning to an alternate state (regime shift, exhibiting early warning signals (EWS)) while the forest-marsh is equivalent to the Slowly Transitioning (scenario 2: ecosystem is transitioning slowly to an alternate state)* |  |  |  |

Furthermore, we compare our results to the annual national land cover dataset, as also done in previous studies (32). We found that the yearly forest loss to marsh from the NLCD data is 165 ha/yr-1, which is about 38.6% lower than our forest-marsh annual loss (269 ha yr-1). It should be noted that we only considered the emergent wetland as a marsh in the NLCD data. Another reason for the large difference could also be due to differences in accuracy, especially given that the overall accuracy of the NLCD 2021 model is 82.82, and the 1985 model is 83.25, with their wetland class overlapping highly with forest (81). In both datasets, the spatial distribution of coastal forest loss within the study area is primarily concentrated along the coast, especially the eastern side of the Alligator River National Wildlife Refuge, exhibiting more significant forest loss compared to the western side of the Refuge (see Fig 6).

Comparing our result to that of Martinez et al. (2024) (62), over the same spatial extent, their annual estimate of Forest-Ghost forest loss is 188 ha yr-1, which is lower than our estimated forest-ghost forest transitions. Furthermore, we compared our results of forest-ghost loss and forest-marsh loss to area loss estimates across all years from 1985 to 2019, as reported by Ury et al (2021) (9). We found that over the same spatial extent, the annual forest loss to marsh is 259 ha yr-1, which is 4% lower than our study estimates. On the other hand, our forest-ghost yearly forest loss is 40% lower than (9) ’s estimates. One possible explanation for this, aside from the 2-year difference between our results, is the comparison across all years rather than between two years. Our higher classification accuracy could also influence the area estimated for each class, ultimately resulting in a difference in total land area.

### 4.3 Drivers of forest change and ghost forest formation

Sea-level rise caused significant forest die-back along the coastal plains of North Carolina. Our time-series land cover maps for 1985, 2010, and 2021 reveal accelerating rates of coastal forest loss, concurrent with the observed rise in regional global sea level over the last 4 decades, as well as climate extremes (32,82). For instance, 23,876 ha of forest were lost between 2010 and 2021 to marsh, ghost forest, and shrub, which is 1.4 times higher than the 16,968 ha lost between 1985 and 2010. Numerous extreme events marked this timeframe (i.e., 2010-2021). The various parts of the study area experienced disturbances to varying degrees, with some occurring alone and others alongside multiple events. Previous studies also (9,62) noted that this area experienced prolonged drought between 2007 and 2011, and Hurricane Irene (both in 2011). While some of these extreme events occurred several years before 2021, some areas never fully recovered, having been pushed beyond their boundary into a new ecosystem state, despite being highly protected.

Specifically, ghost forests had a net gain of 7561 ha (687 ha yr^-1^) between 2010 and 2021 (82% from forest, the remaining 18% primarily from shrub), which is 2.5 times higher than the estimated net gain in ghost forest area between 1985 and 2010 (3087 ha). Additionally, forests are being converted to cultivated land, especially in the western part of our study area, indicating that resilient forest areas could also be subject to persistent anthropogenic land-use changes.

We found the greatest number of ghost forest transitions and expansions of wetlands into coastal forests along the estuary margins and coastlines. In line with Ury et al. (2021) (9), forest loss was concentrated about 1 km inland from the coast margin across our study site. The forest transitions, however, were more extensive along the Alligator River National Wildlife Refuge (Fig 6i), with a greater concentration on the eastern side, indicating a more significant loss of forest compared to the western side. Some forest loss was also observed in the interiors of the study area. One of the major drivers of coastal forest loss, resulting in both ghost forests and marsh, is the proximity to the channel (9), reflecting the vulnerability of coastal forests to inundation and saltwater intrusion. We found that forests lost to marshes, shrubs, and ghost forests were concentrated at an average distance of 1.2 km from major water bodies, though we identified areas of ghost forest formation that appeared up to 17 km away from the coastline and approximately 60km away from the Pamlico Sound, which interconnected drainage networks or heavy storms that result in periods of inundation could have facilitated. This result complements earlier findings in coastal North Carolina (4,9,83).

Similarly, increasing salinity and higher RSLR were positively associated with change, supporting the role of ecological stressors and climate-driven processes such as ghost forest formation. These results are not unexpected, as a similar region along the eastern US coast is already recognized as a global hotspot for accelerated relative sea-level rise (RSLR) (6). For instance, the average RSLRR across all stations in the coastal plain of North Carolina increased from 3.5 mm/year in 2010 to 5.1 mm/year in 2021, indicating a continuous increase in RSLRR from 1985 to 2021 compared to the period from 1985 to 2010 (see S6a Fig). Smart et al. (2020) also reported declines in aboveground biomass and carbon near shorelines and in areas with higher salinities. Our study also demonstrated that these factors are stronger drivers of coastal forest change than elevation and slope alone (4,47). This region has experienced major hurricane events, including Hurricane Irene in 2011, classified as a category 3, which passed through APP as a category 1 (9,62), and Hurricane Julia in 2016, which could contribute to high salinity through inundation.

Interestingly, while elevation significantly affects forest-to-marsh loss, with low-lying relief areas experiencing the greatest forest change, our study reveals that this is not the case for ghost forest formation, indicating that elevation has a weak statistical influence but a larger magnitude on forest conversion to ghost forests. Slope, on the other hand, has a stronger impact in predicting forest-ghost forest compared to forest-marsh. We observed greater forest loss in the downslope area (0-20 degrees, Fig. 7) than on the mid- or upper slope. These areas appear wetter and are highly prone to inundation, while forest loss observed on the upper slope might be attributed to dryness and lag effects of historic drought or fire. Therefore, our observed increased forest loss in downslope areas supports the idea that topography influences gradients in moisture and stress sensitivity (84,85), where increasing slopes tend to reduce the duration of tidal flooding, ultimately decreasing salinization and waterlogging conditions (1,86).

We also note that not only are forests transitioning, but also the conversion of farmland to wetland, which could be related to the abandonment of farmlands in response to salinization facilitated by drainage ditches and other legacy land management practices (45,87). Some of the farmland on the coastal plain of North Carolina has since been transformed into forested wetland ecosystems (88,89). These changes indicate increased flooding and probable salinization of these areas due to sea level rise (8,9,27). We also observed forest regrowth from shrubs, as also reported by Ury et al (2021) (9). Specifically, the patches around the Pocosin Lakes National Wildlife Refuge, which represent a shrub and ghost forests class shown on our 1985 classified map, represent areas that experienced a wildfire event in 1985 (Allan Road Fires, Poulter et al., 2006), which were subsequently replaced by forests in 2021. Some of the ghost forest areas also overlapped with historic wildfire events, specifically the 2011 Pains Bay Fire along the Southern portion of the Alligator Refuge shoreline (4,62). This pattern suggests that multiple mechanisms are contributing to ghost forest formation across the landscape. Some of these mechanisms, i.e., wildfire, are cyclical and include vegetation recovery (62,90). While the spatial extent is relatively small, this highlights the limitations of distinguishing ghost forest cover from other disturbance-related land types or from scanty trees in coarse 30-meter-resolution imagery, given the small training dataset, especially for the pre-2021 images. With our results showing that elevation is not a strong predictor of ghost forest presence and that ghost forest distribution varies widely across elevation gradients, one explanation is that some ditches may serve as pathways for saltwater intrusion, even at higher elevations (18,19). It’s interesting to note that the entire forest loss is relatively concentrated in the low-relief zone (<5 meters, see Fig 7), so the definition of high elevation might have been exaggerated here because even areas considered as “higher elevation” are still low in an absolute sense, with elevation also increasing inland towards the western side of our study. Additionally, potential misclassification between ghost forest and marsh, particularly in low-lying transitional zones, may further obscure the relationship between elevation and ghost forest occurrence. We also observed that some of our forest losses occurred in the interior, indicating that loss is not only concentrated on the shoreline, a result in line with Ury et al (2021) (9)’s findings. Additionally, fire-driven ghost forests may occur more frequently at higher elevations, while saltwater-driven ghost forests occur at lower elevations. This could be explored further in future work by disaggregating ghost forests based on disturbance drivers. Although significant, elevation is not a major predictor of shrub presence, indicating that we observed an expansion of shrubs in high-relief areas of the coastal landscape. However, increased salinity favors shrub growth, consistent with a previous study (91). Their work shows that moderate salinity favors *M. cerifera* growth and expansion, with shrubs advancing to areas that may later become ghost forests or marshes. We also observed that as marshes move inland, they are replacing shrubs (see Fig 6c), which could indicate the development of a shrub-scrub marsh. This could indicate that some of the transitions are happening in areas that are too wet to turn into swamps and too dry or shallow to develop into marshes (92).

## 5 Conclusion

Our analysis underscores the effectiveness of both Landsat and Sentinel imagery in land cover classification, with high overall accuracy and F1 scores for each sensor. Our approach, which utilizes a deep learning algorithm incorporating phenology and multispectral indices, improves the classification accuracy for mapping various coastal forest vegetation types, including ghost forest and shrub, which have underrepresented training samples compared to traditional classifiers employed in previous studies. The results of our study further reveal the extent and spatial patterns of coastal forest retreats, with a striking rate of transition over 37 years, matching existing remote sensing-based products (e.g., NLCD). Furthermore, this study reveals that proximity to channels, high salinity, low elevation, shallow slope, and rapid relative sea level rise (RSLR) influence the conversion of forest to other land cover differently. The relative intensification of climate change, characterized by observed increases in sea level rise and maximum temperature, will continue to threaten coastal ecosystems, further exacerbated by enhanced saltwater intrusion through channels and ditches, as well as extreme events. Continuous documentation of emerging ghost forests and shrub growth is crucial for training models to enhance our understanding and knowledge of coastal ecosystems’ transgressions. Information about the rate of transgressions could serve as a valuable resource for targeted restoration and reconstruction efforts that might help return some disturbed areas to their near-original state and prevent new areas from transitioning.

## Acknowledgements

This research is funded through the Southeast Climate Adaptation Science Centers (SECASC) Global Change Research fellowship (Grant Number: G23AC00548). We extend our gratitude to Dr. Ury for providing some training data to support this work.

## Supporting Information

**Figure S1.** Description of extracted phenological metrics from the NDVI

**Figure S2.** (a) Landsat 8 model’s loss curve, and (b) Sentinel 2 model’s loss curve

**Figure S3.** Permuted variable contributions to classification model performance for each vegetation type in the Landsat 8 full-feature CNN classification model. Bars with positive values indicate statistically significant predictors (those on the negative side indicate the predictor is not statistically significant). The X-axis represents the mean decrease in accuracy.

**Figure S4.** Permuted variable contributions to classification model performance for each vegetation type in the Sentinel Full features CNN classification model. Bars with positive values indicate statistically significant predictors (those on the negative side indicate the predictor is not statistically significant). The X-axis represents the mean decrease in accuracy.

**Figure S5.** Permuted variable feature importance for the non-phenology model for (a) Landsat 8 and (b) Sentinel 2

**Figure S6.** (a) Relative sea level rise, and (b) Distribution of vegetation samples along elevation for the years 1985, 2010, and 2021.

**Figure S7.** Mean EVI among forest change pathways for 1985 and 2021.

## References

1. Chen Y, Kirwan ML. Upland forest retreat lags behind sea-level rise in the MID-ATLANTIC coast. Global Change Biology. 2024 Jan;30(1):e17081. doi:10.1111/gcb.17081

2. Kirwan ML, Gedan KB. Sea-level driven land conversion and the formation of ghost forests. Nat Clim Chang. 2019 Jun;9(6):6. doi:10.1038/s41558-019-0488-7

3. McDermott A. Ghost forests haunt the East Coast, harbingers of sea-level rise. Proc Natl Acad Sci USA. 2023 Sep 19;120(38):e2314607120. doi:10.1073/pnas.2314607120

4. Smart LS, Taillie PJ, Poulter B, Vukomanovic J, Singh KK, Swenson JJ, et al. Aboveground carbon loss associated with the spread of ghost forests as sea levels rise. Environ Res Lett. 2020 Oct 1;15(10):10. doi:10.1088/1748-9326/aba136

5. Osland MJ, Chivoiu B, Enwright NM, Thorne KM, Guntenspergen GR, Grace JB, et al. Migration and transformation of coastal wetlands in response to rising seas. Sci Adv. 2022 Jul;8(26):eabo5174. doi:10.1126/sciadv.abo5174

6. Sallenger AH, Doran KS, Howd PA. Hotspot of accelerated sea-level rise on the Atlantic coast of North America. Nature Clim Change. 2012 Dec;2(12):884–8. doi:10.1038/nclimate1597

7. White E, Ury EA, Bernhardt ES, Yang X. Climate Change Driving Widespread Loss of Coastal Forested Wetlands Throughout the North American Coastal Plain. Ecosystems. 2022 Jun;25(4):812–27. doi:10.1007/s10021-021-00686-w

8. Schieder NW, Kirwan ML. Sea-level driven acceleration in coastal forest retreat. Geology. 2019 Dec 1;47(12):1151–5. doi:10.1130/G46607.1

9. Ury EA, Yang X, Wright JP, Bernhardt ES. Rapid deforestation of a coastal landscape driven by sea-level rise and extreme events. Ecological Applications. 2021 Jul;31(5):5. doi:10.1002/eap.2339

10. Grieger R, Capon SJ, Hadwen WL, Mackey B. Between a bog and a hard place: a global review of climate change effects on coastal freshwater wetlands. Climatic Change. 2020 Nov;163(1):161–79. doi:10.1007/s10584-020-02815-1

11. Ury EA, Anderson SM, Peet RK, Bernhardt ES, Wright JP. Succession, regression and loss: does evidence of saltwater exposure explain recent changes in the tree communities of North Carolina’s Coastal Plain? Annals of Botany. 2019 Apr 6. doi:10.1093/aob/mcz039

12. Williams K, Ewel KC, Stumpf RP, Putz FE, Workman TW. SEA-LEVEL RISE AND COASTAL FOREST RETREAT ON THE WEST COAST OF FLORIDA, USA. Ecology. 1999 Sep;80(6):2045–63. doi:10.1890/0012-9658(1999)080[2045:SLRACF]2.0.CO;2

13. Brinson MM, Christian RR, Blum LK. Multiple States in the Sea-Level Induced Transition from Terrestrial Forest to Estuary. Estuaries. 1995 Dec;18(4):4. doi:10.2307/1352383

14. Ardón M, Potter KM, White E, Woodall CW. Coastal carbon sentinels: A decade of forest change along the eastern shore of the US signals complex climate change dynamics. Ashraf MI, editor. PLOS Clim. 2025 Jan 9;4(1):e0000444. doi:10.1371/journal.pclm.0000444

15. Bhattachan A, Emanuel RE, Ardón M, Bernhardt ES, Anderson SM, Stillwagon MG, et al. Evaluating the effects of land-use change and future climate change on vulnerability of coastal landscapes to saltwater intrusion. Zak DR, Olden JD, editors. Elementa: Science of the Anthropocene. 2018 Jan 1;6:62. doi:10.1525/elementa.316

16. Herbert ER, Boon P, Burgin AJ, Neubauer SC, Franklin RB, Ardón M, et al. A global perspective on wetland salinization: ecological consequences of a growing threat to freshwater wetlands. Ecosphere. 2015 Oct;6(10):1–43. doi:10.1890/ES14-00534.1

17. White E, Kaplan D. Restore or retreat? saltwater intrusion and water management in coastal wetlands. Ecosyst Health Sustain. 2017 Jan;3(1):e01258. doi:10.1002/ehs2.1258

18. Ardón M, Morse JL, Colman BP, Bernhardt ES. Drought-induced saltwater incursion leads to increased wetland nitrogen export. Global Change Biology. 2013 Oct;19(10):2976–85. doi:10.1111/gcb.12287

19. Neville JA, Emanuel RE, Ardón M, Pavelsky T. Location and Design of Flow Control Structures Differentially Influence Salinity Patterns in Small Artificial Drainage Systems. J Water Resour Plann Manage. 2023 Jun;149(6):6. doi:10.1061/JWRMD5.WRENG-5840

20. O’Donnell KL, Bernhardt ES, Yang X, Emanuel RE, Ardón M, Lerdau MT, et al. Saltwater intrusion and sea level rise threatens U.S. rural coastal landscapes and communities. Anthropocene. 2024 Mar;45:100427. doi:10.1016/j.ancene.2024.100427

21. Berner LT, Goetz SJ. Satellite observations document trends consistent with a boreal forest biome shift. Global Change Biology. 2022 May;28(10):3275–92. doi:10.1111/gcb.16121

22. Decuyper M, Chávez RO, Lohbeck M, Lastra JA, Tsendbazar N, Hackländer J, et al. Continuous monitoring of forest change dynamics with satellite time series. Remote Sensing of Environment. 2022 Feb;269:112829. doi:10.1016/j.rse.2021.112829

23. Pu R, Landry S, Yu Q. Assessing the potential of multi-seasonal high resolution Pléiades satellite imagery for mapping urban tree species. International Journal of Applied Earth Observation and Geoinformation. 2018 Sep;71:144–58. doi:10.1016/j.jag.2018.05.005

24. Sexton JO, Urban DL, Donohue MJ, Song C. Long-term land cover dynamics by multi-temporal classification across the Landsat-5 record. Remote Sensing of Environment. 2013 Jan;128:246–58. doi:10.1016/j.rse.2012.10.010

25. De Beurs KM, Henebry GM. Spatio-Temporal Statistical Methods for Modelling Land Surface Phenology. In: Hudson IL, Keatley MR, editors. Phenological Research [Internet]. Dordrecht: Springer Netherlands; 2010 [cited 2025 Oct 14]. p. 177–208. Available from: http://link.springer.com/10.1007/978-90-481-3335-2_9doi:10.1007/978-90-481-3335-2_9

26. Reed BC, Brown JF, VanderZee D, Loveland TR, Merchant JW, Ohlen DO. Measuring phenological variability from satellite imagery. J Vegetation Science. 1994 Oct;5(5):703–14. doi:10.2307/3235884

27. Gray PC, Chamorro DF, Ridge JT, Kerner HR, Ury EA, Johnston DW. Temporally Generalizable Land Cover Classification: A Recurrent Convolutional Neural Network Unveils Major Coastal Change through Time. Remote Sensing. 2021 Oct 2;13(19):19. doi:10.3390/rs13193953

28. Hughes L. Biological consequences of global warming: is the signal already apparent? Trends in Ecology & Evolution. 2000 Feb;15(2):56–61. doi:10.1016/S0169-5347(99)01764-4

29. Scranton K, Amarasekare P. Predicting phenological shifts in a changing climate. Proc Natl Acad Sci USA. 2017 Dec 12;114(50):13212–7. doi:10.1073/pnas.1711221114

30. Thackeray SJ, Sparks TH, Frederiksen M, Burthe S, Bacon PJ, Bell JR, et al. Trophic level asynchrony in rates of phenological change for marine, freshwater and terrestrial environments. Global Change Biology. 2010 Dec;16(12):3304–13. doi:10.1111/j.1365-2486.2010.02165.x

31. Chang T, Rasmussen B, Dickson B, Zachmann L. Chimera: A Multi-Task Recurrent Convolutional Neural Network for Forest Classification and Structural Estimation. Remote Sensing. 2019 Mar 29;11(7):768. doi:10.3390/rs11070768

32. Chen Y, Kirwan ML. A phenology- and trend-based approach for accurate mapping of sea-level driven coastal forest retreat. Remote Sensing of Environment. 2022 Nov;281:113229. doi:10.1016/j.rse.2022.113229

33. Diao C, Wang L. Incorporating plant phenological trajectory in exotic saltcedar detection with monthly time series of Landsat imagery. Remote Sensing of Environment. 2016 Sep;182:60–71. doi:10.1016/j.rse.2016.04.029

34. Mountrakis G, Im J, Ogole C. Support vector machines in remote sensing: A review. ISPRS Journal of Photogrammetry and Remote Sensing. 2011 May;66(3):247–59. doi:10.1016/j.isprsjprs.2010.11.001

35. Belgiu M, Drăguţ L. Random forest in remote sensing: A review of applications and future directions. ISPRS Journal of Photogrammetry and Remote Sensing. 2016 Apr;114:24–31. doi:10.1016/j.isprsjprs.2016.01.011

36. Cheng G, Yang C, Yao X, Guo L, Han J. When Deep Learning Meets Metric Learning: Remote Sensing Image Scene Classification via Learning Discriminative CNNs. IEEE Trans Geosci Remote Sensing. 2018 May;56(5):2811–21. doi:10.1109/TGRS.2017.2783902

37. Gómez C, White JC, Wulder MA. Optical remotely sensed time series data for land cover classification: A review. ISPRS Journal of Photogrammetry and Remote Sensing. 2016 Jun;116:55–72. doi:10.1016/j.isprsjprs.2016.03.008

38. Ma L, Liu Y, Zhang X, Ye Y, Yin G, Johnson BA. Deep learning in remote sensing applications: A meta-analysis and review. ISPRS Journal of Photogrammetry and Remote Sensing. 2019 Jun;152:166–77. doi:10.1016/j.isprsjprs.2019.04.015

39. Madonsela S, Cho MA, Mathieu R, Mutanga O, Ramoelo A, Kaszta Ż, et al. Multi-phenology WorldView-2 imagery improves remote sensing of savannah tree species. International Journal of Applied Earth Observation and Geoinformation. 2017 Jun;58:65–73. doi:10.1016/j.jag.2017.01.018

40. Liu Y, Chen X, Wang Z, Wang ZJ, Ward RK, Wang X. Deep learning for pixel-level image fusion: Recent advances and future prospects. Information Fusion. 2018 Jul;42:158–73. doi:10.1016/j.inffus.2017.10.007

41. Zhong Y, Han X, Zhang L. Multi-class geospatial object detection based on a position-sensitive balancing framework for high spatial resolution remote sensing imagery. ISPRS Journal of Photogrammetry and Remote Sensing. 2018 Apr;138:281–94. doi:10.1016/j.isprsjprs.2018.02.014

42. Schmidhuber J. Deep learning in neural networks: An overview. Neural Networks. 2015 Jan;61:85–117. doi:10.1016/j.neunet.2014.09.003

43. Lecun Y, Bottou L, Bengio Y, Haffner P. Gradient-based learning applied to document recognition. Proc IEEE. 1998 Nov;86(11):2278–324. doi:10.1109/5.726791

44. Razzaghi P, Abbasi K, Bayat P. Learning spatial hierarchies of high-level features in deep neural network. Journal of Visual Communication and Image Representation. 2020 Jul;70:102817. doi:10.1016/j.jvcir.2020.102817

45. Thomas VA, Wynne RH, Kauffman J, McCurdy W, Brooks EB, Thomas RQ, et al. Mapping thins to identify active forest management in southern pine plantations using Landsat time series stacks. Remote Sensing of Environment. 2021 Jan;252:112127. doi:10.1016/j.rse.2020.112127

46. Martinez M, Ardón M. Drivers of greenhouse gas emissions from standing dead trees in ghost forests. Biogeochemistry. 2021 Jul;154(3):471–88. doi:10.1007/s10533-021-00797-5

47. Taillie PJ, Moorman CE, Poulter B, Ardón M, Emanuel RE. Decadal-Scale Vegetation Change Driven by Salinity at Leading Edge of Rising Sea Level. Ecosystems. 2019 Dec;22(8):1918–30. doi:10.1007/s10021-019-00382-w

48. U.S. Geological Survey (USGS) Gap Analysis Project (GAP). Protected Areas Database of the United States (PAD-US) 4 [Internet]. U.S. Geological Survey; [cited 2026 Apr 9]. Available from: https://www.sciencebase.gov/catalog/item/65294599d34e44db0e2ed7cfdoi:10.5066/P96WBCHS

49. Richardson CJ. Pocosins: Hydrologically isolated or integrated wetlands on the landscape? Wetlands. 2003 Sep;23(3):563–76. doi:10.1672/0277-5212(2003)023[0563:PHIOIW]2.0.CO;2

50. Earth Resources Observation and Science (EROS) Center. Landsat 8-9 Operational Land Imager / Thermal Infrared Sensor Level-2, Collection 2 [dataset] [Internet]. U.S. Geological Survey: 10.5066/P9OGBGM6; 2020. Available from: 10.5066/P9OGBGM6

51. Wendelberger K, Gann D, Richards J. Using Bi-Seasonal WorldView-2 Multi-Spectral Data and Supervised Random Forest Classification to Map Coastal Plant Communities in Everglades National Park. Sensors. 2018 Mar 9;18(3):829. doi:10.3390/s18030829

52. NC OneMap. NC Orthoimagery Program, North Carolina Department of Information Technology, Government Data Analytics Center, Center for Geographic Information and Analysis. [Internet]. 2020 [cited 2026 Mar 2]. Available from: https://services.nconemap.gov/secure/rest/services/Imagery

53. National Agriculture Imagery Program (NAIP). USGS Earth Resources Observation and Science (EROS) Center Archive - Aerial Photography - [Internet]. 2019 [cited 2024 Dec 1]. Available from: https://www.usgs.gov/centers/eros/science/usgs-eros-archive-aerial-photography-national-agriculture-imagery-program-naip

54. Zeng L, Wardlow BD, Xiang D, Hu S, Li D. A review of vegetation phenological metrics extraction using time-series, multispectral satellite data. Remote Sensing of Environment. 2020 Feb;237:111511. doi:10.1016/j.rse.2019.111511

55. Caparros-Santiago JA, Rodriguez-Galiano V, Dash J. Land surface phenology as indicator of global terrestrial ecosystem dynamics: A systematic review. ISPRS Journal of Photogrammetry and Remote Sensing. 2021 Jan;171:330–47. doi:10.1016/j.isprsjprs.2020.11.019

56. Savitzky Abraham, Golay MJE. Smoothing and Differentiation of Data by Simplified Least Squares Procedures. Anal Chem. 1964 Jul 1;36(8):1627–39. doi:10.1021/ac60214a047

57. Ivits E, Horion S, Fensholt R, Cherlet M. Drought footprint on E uropean ecosystems between 1999 and 2010 assessed by remotely sensed vegetation phenology and productivity. Global Change Biology. 2014 Feb;20(2):581–93. doi:10.1111/gcb.12393

58. Shen M, Piao S, Cong N, Zhang G, Jassens IA. Precipitation impacts on vegetation spring phenology on the T ibetan P lateau. Global Change Biology. 2015 Oct;21(10):3647–56. doi:10.1111/gcb.12961

59. Lewis T. [Internet]. 2020. Available from: https://github.com/lewistrotter/PhenoloPy

60. Zhang L, Shen M, Liu L, Chen X, Cao R, Dong Q, et al. Refining landsat-based annual NDVImax estimation using shape model fitting and phenological metrics. Ecological Informatics. 2025 Jul;87:103107. doi:10.1016/j.ecoinf.2025.103107

61. Kruskal WH, Wallis WA. Use of Ranks in One-Criterion Variance Analysis. Journal of the American Statistical Association. 1952 Dec;47(260):583–621. doi:10.1080/01621459.1952.10483441

62. Martinez M, Ardón M, Gray J. Detecting Trajectories of Regime Shifts and Loss of Resilience in Coastal Wetlands using Remote Sensing. Ecosystems. 2024 Dec;27(8):1060–75. doi:10.1007/s10021-024-00938-5

63. Healey S, Cohen W, Zhiqiang Y, Krankina O. Comparison of Tasseled Cap-based Landsat data structures for use in forest disturbance detection. Remote Sensing of Environment. 2005 Aug 15;97(3):301–10. doi:10.1016/j.rse.2005.05.009

64. Baig MHA, Zhang L, Shuai T, Tong Q. Derivation of a tasselled cap transformation based on Landsat 8 at-satellite reflectance. Remote Sensing Letters. 2014 May 4;5(5):423–31. doi:10.1080/2150704X.2014.915434

65. Nedkov R. Orthogonal transformation of segmented images from the satellite sentinel-2. 2017,70, 687–692. Proceedings of the Bulgarian Academy of Sciences. 70(5):687–92.

66. Danielson JJ, Poppenga SK, Tyler DJ, Palaseanu-Lovejoy M, Gesch DB. Coastal National Elevation Database: U.S. Geological Survey Fact Sheet [Fact Sheet]. 2018. (Fact Sheet).

67. Phiri D, Morgenroth J. Developments in Landsat Land Cover Classification Methods: A Review. Remote Sensing. 2017 Sep 19;9(9):967. doi:10.3390/rs9090967

68. Fieres J, Schemmel J, Meier K. Training convolutional networks of threshold neurons suited for low-power hardware implementation. In: The 2006 IEEE International Joint Conference on Neural Network Proceedings [Internet]. Vancouver, BC, Canada: IEEE; 2006 [cited 2025 Jun 21]. p. 21–8. Available from: http://ieeexplore.ieee.org/document/1716065/doi:10.1109/IJCNN.2006.246654

69. Indolia S, Goswami AK, Mishra SP, Asopa P. Conceptual Understanding of Convolutional Neural Network-A Deep Learning Approach. Procedia Computer Science. 2018;132:679–88. doi:10.1016/j.procs.2018.05.069

70. LeCun Y, Bengio Y, Hinton G. Deep learning. Nature. 2015 May 28;521(7553):436–44. doi:10.1038/nature14539

71. Maggiori E, Tarabalka Y, Charpiat G, Alliez P. Convolutional Neural Networks for Large-Scale Remote-Sensing Image Classification. IEEE Trans Geosci Remote Sensing. 2017 Feb;55(2):645–57. doi:10.1109/TGRS.2016.2612821

72. Wang S, Chen W, Xie SM, Azzari G, Lobell DB. Weakly Supervised Deep Learning for Segmentation of Remote Sensing Imagery. Remote Sensing. 2020 Jan 7;12(2):207. doi:10.3390/rs12020207

73. Lee KB, Cheon S, Kim CO. A Convolutional Neural Network for Fault Classification and Diagnosis in Semiconductor Manufacturing Processes. IEEE Trans Semicond Manufact. 2017 May;30(2):135–42. doi:10.1109/TSM.2017.2676245

74. Palsson F, Sveinsson JR, Ulfarsson MO. Multispectral and Hyperspectral Image Fusion Using a 3-D-Convolutional Neural Network. IEEE Geosci Remote Sensing Lett. 2017 May;14(5):639–43. doi:10.1109/LGRS.2017.2668299

75. Nicolau AP, Dyson K, Saah D, Clinton N. Accuracy Assessment: Quantifying Classification Quality. In: Cardille JA, Crowley MA, Saah D, Clinton NE, editors. Cloud-Based Remote Sensing with Google Earth Engine [Internet]. Cham: Springer International Publishing; 2024 [cited 2024 Dec 1]. p. 135–45. Available from: https://link.springer.com/10.1007/978-3-031-26588-4_7doi:10.1007/978-3-031-26588-4_7

76. Mandler H, Weigand B. Feature importance in neural networks as a means of interpretation for data-driven turbulence models. Computers & Fluids. 2023 Oct;265:105993. doi:10.1016/j.compfluid.2023.105993

77. Cheon W, Han M, Jeong S, Oh ES, Lee SU, Lee SB, et al. Feature Importance Analysis of a Deep Learning Model for Predicting Late Bladder Toxicity Occurrence in Uterine Cervical Cancer Patients. Cancers. 2023 Jul 2;15(13):3463. doi:10.3390/cancers15133463

78. The Multi-Resolution Land Characteristics (MRLC) consortium. National Land Cover Database Class Legend and Description [Internet]. 2025. Available from: https://www.mrlc.gov/data/legends/national-land-cover-database-class-legend-and-description

79. Emadi M, Baghernejad M. Comparison of spatial interpolation techniques for mapping soil pH and salinity in agricultural coastal areas, northern Iran. Archives of Agronomy and Soil Science. 2014 Sep 2;60(9):1315–27. doi:10.1080/03650340.2014.880837

80. Sambandham VT, Kirchheim K, Ortmeier F, Mukhopadhaya S. Deep learning-based harmonization and super-resolution of Landsat-8 and Sentinel-2 images. ISPRS Journal of Photogrammetry and Remote Sensing. 2024 Jun;212:274–88. doi:10.1016/j.isprsjprs.2024.04.026

81. United States Geological Survey. Annual National Land Cover Database (NLCD) Collection 1.0 Validation Tables [Zip,jpg,csv,xml,pdf]. U.S. Geological Survey; 2025 [cited 2025 Sep 17]. Available from: https://www.sciencebase.gov/catalog/item/6813a741d4be02316305177ddoi:10.5066/P1KJXXGA

82. Dangendorf S, Marcos M, Wöppelmann G, Conrad CP, Frederikse T, Riva R. Reassessment of 20th century global mean sea level rise. Proc Natl Acad Sci USA. 2017 Jun 6;114(23):5946–51. doi:10.1073/pnas.1616007114

83. Poulter B, Goodall JL, Halpin PN. Applications of network analysis for adaptive management of artificial drainage systems in landscapes vulnerable to sea level rise. Journal of Hydrology. 2008 Aug;357(3–4):207–17. doi:10.1016/j.jhydrol.2008.05.022

84. Elliott KJ, Vose JM, Swank WT, Bolstad PV. Long-Term Patterns in Vegetation-Site Relationships in a Southern Appalachian Forest. Journal of the Torrey Botanical Society. 1999 Oct;126(4):320. doi:10.2307/2997316

85. Hawthorne S, Miniat CF. Topography may mitigate drought effects on vegetation along a hillslope gradient. Ecohydrology. 2018 Jan;11(1):e1825. doi:10.1002/eco.1825

86. Hussein AH, Rabenhorst MC. Tidal Inundation of Transgressive Coastal Areas: Pedogenesis of Salinization and Alkalinization. Soil Science Soc of Amer J. 2001 Mar;65(2):536–44. doi:10.2136/sssaj2001.652536x

87. Tully K, Gedan K, Epanchin-Niell R, Strong A, Bernhardt ES, Bendor T, et al. Corrigendum: The Invisible Flood: The Chemistry, Ecology, and Social Implications of Coastal Saltwater Intrusion. BioScience. 2019 Sep 1;69(9):760–760. doi:10.1093/biosci/biz083

88. USDA Natural Resources Conservation Service (NRCS). Partnership Powers Large-Scale Restoration: How NRCS and Partners Transform North Carolina’s Coastal Landscape [Internet]. [cited 2026 Apr 1]. Available from: https://www.nrcs.usda.gov/state-offices/north-carolina/news/partnership-powers-large-scale-restoration-how-nrcs-and-partners

89. Ardón M, Montanari S, Morse JL, Doyle MW, Bernhardt ES. Phosphorus export from a restored wetland ecosystem in response to natural and experimental hydrologic fluctuations. J Geophys Res. 2010 Dec;115(G4):2009JG001169. doi:10.1029/2009JG001169

90. Poulter B, Christensen NL, Halpin PN. Carbon emissions from a temperate peat fire and its relevance to interannual variability of trace atmospheric greenhouse gases. J Geophys Res. 2006 Mar 27;111(D6):2005JD006455. doi:10.1029/2005JD006455

91. Sward R, Philbrick A, Morreale J, Baird CJ, Gedan K. Shrub expansion in maritime forest responding to sea level rise. Front For Glob Change. 2023 May 5;6:1167880. doi:10.3389/ffgc.2023.1167880

92. Kalliola R, Maarit P, Salo J, Tuomisto H, Ruokolainen K. The Dynamics, Distribution and Classification of Swamp Vegetation in Peruvian Amazonia. Annales Botanici Fennici. 1991;28(3):225–339.

